# Bioaugmentation in anaerobic digesters: A systematic review

**DOI:** 10.1101/2025.01.22.634285

**Authors:** Mozhdeh Alipoursarbani, Jeroen Tideman, Mitzy López, Christian Abendroth

## Abstract

Bioaugmentation, the intentional introduction of specific microorganisms into anaerobic digestion (AD) systems, has shown promise in enhancing methane production and in mitigating stressful conditions, particularly in systems operating below optimal performance. This review presents a systematic literature review (SLR) and meta-analysis to evaluate the efficacy of bioaugmentation strategies in AD. This review identified and analyzed studies meeting predefined eligibility criteria through a structured methodology involving research protocol, search, appraisal, synthesis, analysis, and reporting. A notable innovation of this review is its comprehensive critical comparison of different controls used in bioaugmentation studies, which has been inadequately addressed in previous literature. To facilitate the functional understanding, strains for bioaugmentation were grouped into the four phases of anaerobic digestion (hydrolysis, acidogenesis, acetogenesis and methanogenesis). A highly diverse set of microbes has been described for bioaugmentation, especially from the families *Clostridiaceae*, *Pseudomonadaceae* and *Syntrophomonadaceae*. Most works are related to hydrolysis. The few works that address acidogenesis are mostly related to dark fermentation. Several studies used methanogenic archaea as well as syntrophic acetate oxidising bacteria, despite the difficulties in culturing them. On the other hand, studies applying strains for acetogenesis were largely underrepresented. Especially works on syntrophic propionate and butyrate oxidation (SPO and SBO) were missing.

## 1. Introduction

Fossil fuels continue to play a central role in supplying energy for both industrial and domestic purposes. However, this fuel presents two significant issues: its finite nature leading to eventual depletion and its environmental impact through greenhouse gas emissions, contributing to global warming. Thus, the development of renewable and environmentally friendly alternative energies is crucial to address these limitations.

In recent years, there has been an increasing trend towards the utilisation of anaerobic digestion (AD) for the production of biogas and renewable energy. AD offers a sustainable and environmentally friendly solution for the treatment of organic waste and the generation of valuable resources (Tian et al., 2019). The utilisation of AD to convert organic waste into biogas is a highly effective approach to waste management (Liu et al., 2023b). AD is widely acknowledged as a technology for extracting energy (CH_4_) from organic waste through the action of various microbial communities within anaerobic conditions (Im et al., 2020). This process encompasses several stages including hydrolysis, acidogenesis, acetogenesis, and methanogenesis, with each phase being facilitated by specific microbial consortia (Li et al., 2021). The efficiency of AD depends on the metabolic activities and interactions of the microorganisms (Xu et al., 2023).

The alteration in microbial balance within bioreactors, often caused by the inhibition of certain groups of microorganism or the proliferation of others, is predominantly triggered by various inhibitory factors. These factors encompass elevated levels of inorganic toxicants like ammonium, phosphate, sulfate, and metal ions. Additionally, fluctuations in parameters such as temperature, pH, organic loading rate (OLR), and the resistance of feedstock to biodegradation contribute to decreased efficiency in AD. Among the proposed mitigation strategies are feedstock pretreatment (including ensilage) or dilution, implementation of multi-phase bioreactors, and precise control of temperature and pH, among others. While these strategies show promise, they may also extend the duration and consequently increase the cost of the AD process (Jain et al., 2015).

Given that the factors mentioned above induce shifts in microbial community dynamics, bioaugmentation emerges as a potential alternative strategy to address these limitations. In regard to anaerobic digestion, bioaugmentation involves the introduction of specific stress-resistant or efficient microorganisms into the underlying microbial community with the aim of bolstering its capacity to produce biomethane. This approach has demonstrated success in aerobic biodegradation scenarios, particularly in soil and wastewater, targeting contaminants typically resistant to degradation (Nzila et al., 2016; Semrany et al., 2012; Tyagi et al., 2011).

However, despite the advancements in AD technology, there are still significant challenges to overcome, particularly in the area of biodegradation enhancement or bioaugmentation. Bioaugmentation involves the addition of specific microbial consortia or enzymes to improve the breakdown of complex organic substrates and enhance the biogas production process. The successful implementation of bioaugmentation techniques holds great potential for optimising biogas yields, improving process stability / robustness, and facilitating the digestion of challenging feedstocks (Zhang et al., 2018). This literature review aims to explore the current state of AD for biogas production, with a specific focus on the challenges associated with bioaugmentation.

## 2. Material and methods

### 2.1 Data collection

This review follows a systematic literature review (SLR) approach inspired by, Mengist et al. (2020) aiming to ensure thoroughness, transparency, and reproducibility in identifying and analyzing relevant literature. The systematic process utilised for conducting the search for relevant literature is illustrated in Fig. 1. The authors conducted a comprehensive examination of Clarivate’s Web of Science (WoS) core collection to identify all publications related to the subject of canaerobic OR biogas AND bio$augmentation). The search was performed on December 4, 2022, encompassing a period of 25 years from January 1, 1999, to March 31, 2024. A total of 1058 documents were retrieved from the search. The parsing and analysis of the WoS corpus were carried out using the bibliometrix package in R. Subsequently, 130 review articles were excluded from the analysis, resulting in 928 articles for further examination. Among these articles, 293 specifically focused on bioaugmentation and included defined taxonomic affiliations. Within this subset of bioaugmentation papers, 89 articles specifically addressed bioaugmentation in the context of anaerobic digestion, with taxonomic affiliations also being defined. Of these articles, some could not be accessed, so only the abstracts were evaluated. In total, 635 articles were excluded due to the lack of taxonomic affiliations, with a subset of these focusing on anaerobic digestion.

**Figure 1.**
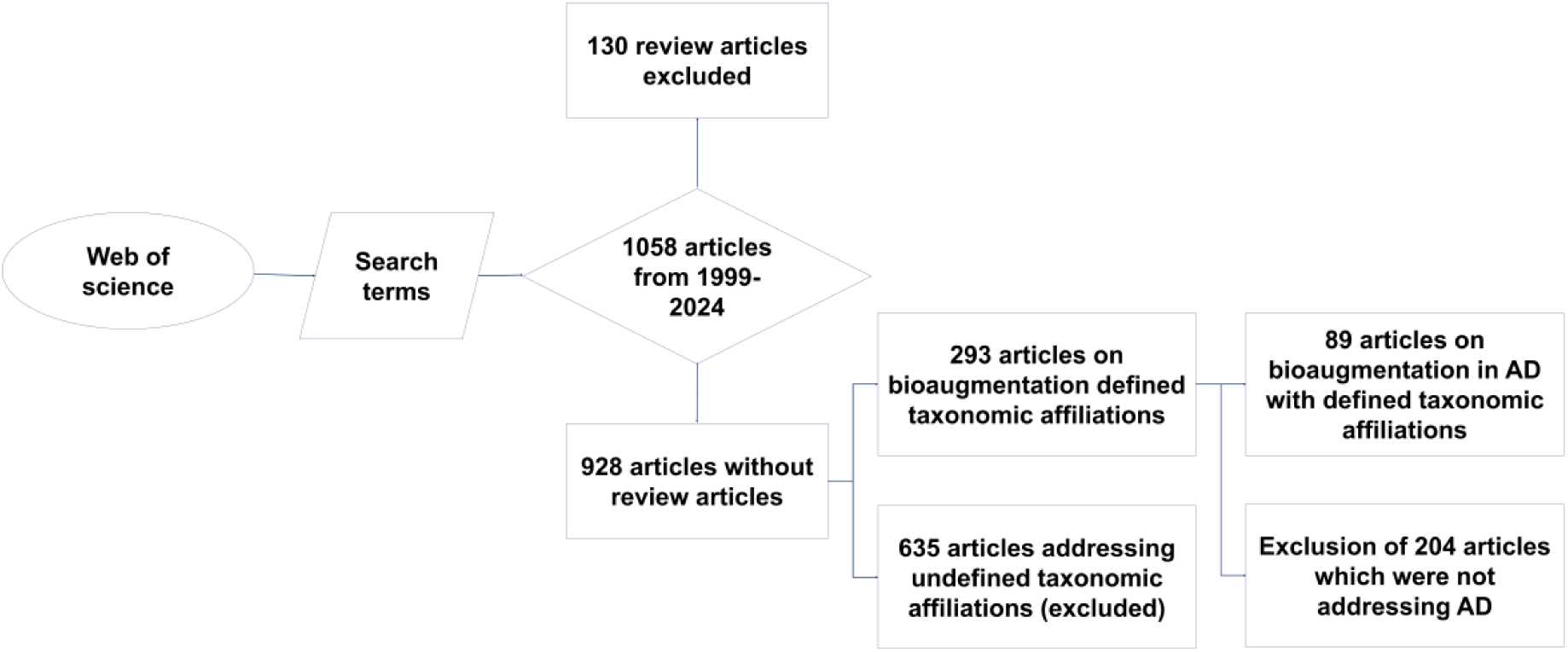
Flow chart depicting the methodology for conducting a systematic review by using Web of Science with the search terms “anaerobic OR biogas AND bio$augmentation”.

## 3. Results and discussion

### 3.1 Exclusion criteria

The distribution of various types of articles are displayed in Fig. 2 (a). The predominant type of article identified was research papers. This finding suggests that researchers have dedicated significant efforts to investigating and contributing new knowledge in the subject area. Although bioaugmentation in the context of anaerobic digestion is a young field of research (beginning in 1999), there is already a considerable amount of review articles (130). Web of Science groups several articles into “Other” and “Meetings”. Considering that after applying the exclusion criteria, relatively few articles remained, it was decided to also take into account “Other” articles and as well articles from “Meetings”. Recently, it has been highlighted as a problem that many review articles are currently reviewing other review articles (Kirchherr, 2023). In agreement with this finding, all review articles were excluded from the reviewed set of articles. Nevertheless, the article should be differentiated from previous review articles in order to better emphasise the importance of the article. This has been detailed in in section 3.2.

**Figure 2.**
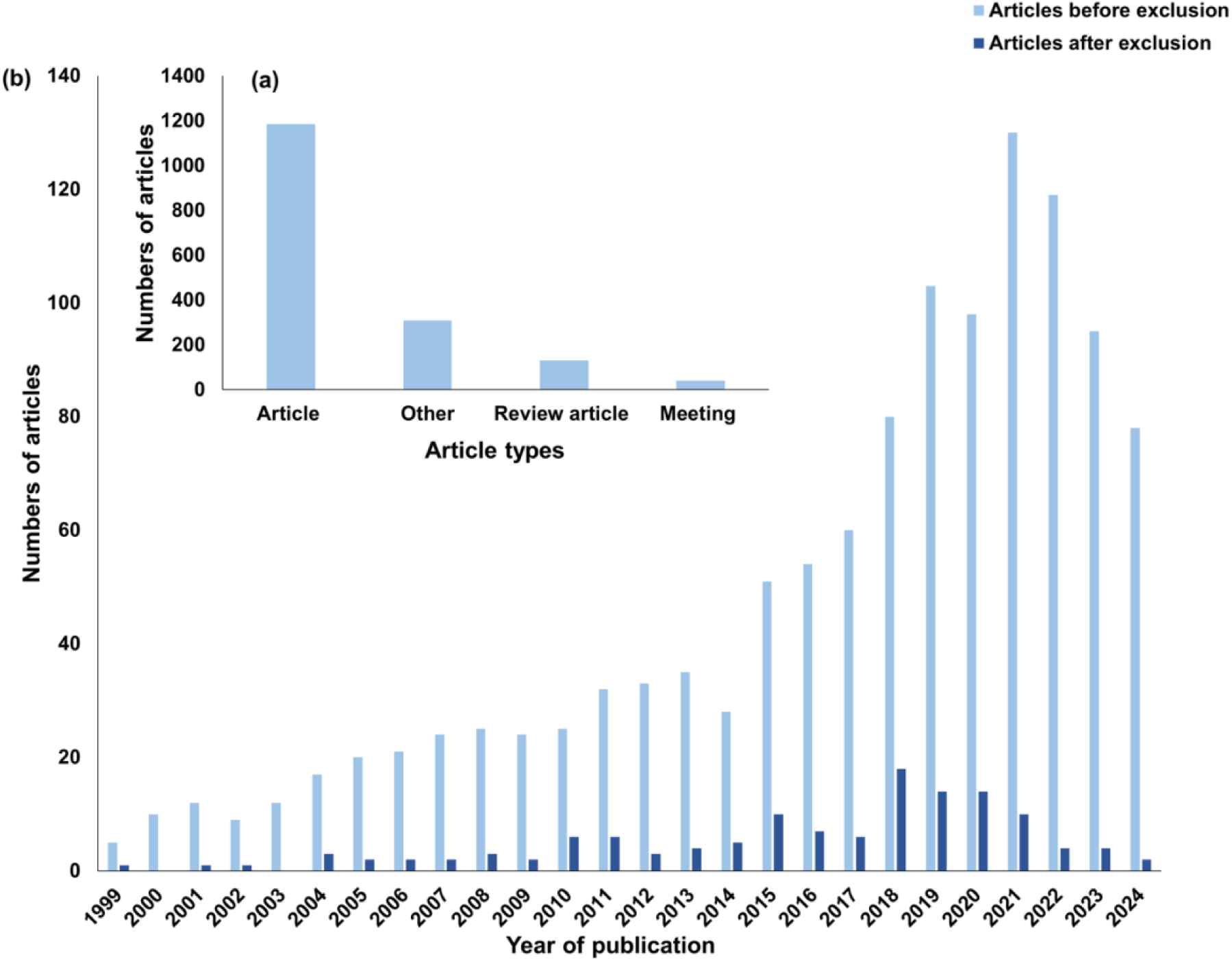
Article types and number of publications: Article types of raw data (a); number of publications per year before and after the application of exclusion criteria (b).

Beginning with a few items in 1999 and the early 2000s, the number of items increased and now indicates exponential growth (Fig. 2b). This trend indicates a significant increase in the attention and interest of the scientific community towards this topic.

Important keywords, their frequency, important journals as well as geographical distribution is shown in Fig. 3. Analysing the occurrence of important terms, it appears that the focus of interest was shifting over the years (Fig. 3a). The terms are ranked by frequency on the right side, reflecting their prominence in the literature. The connections between terms show the evolving research focus over time, revealing how topics have shifted as the field has developed. The early phase of research, represented by the leftmost terms, shows a strong focus on chemical processes involving chlorinated compounds. Terms such as “*dehalogenation*,” “*tetrachloroethene*,” “*chlorinated ethenes*,” “*reductive dechlorination*,” and “*vinyl chloride*” appear frequently and are closely connected. This suggests that early studies were primarily concerned with understanding and mitigating the environmental and health impacts of specific chlorinated organic pollutants. As research progressed, there was a noticeable shift towards microbial processes and biodegradation mechanisms. Terms such as “*culture*,” “*biodegradation*,” and “*reduction*” begin to appear, indicating that researchers started focusing on biological approaches for breaking down these pollutants. This shift reflects an increased interest in utilizing natural microbial communities and bioaugmentation to enhance pollutant degradation, which is evident from the term *bioaugmentation*, one of the most frequently occurring topics in the graph. In more recent years, the focus has expanded to encompass broader environmental applications and sustainable waste management practices. Terms like “*methane production*,” “*co-digestion*,” “*food waste*,” and “*stability*” suggest a growing emphasis on integrating waste management practices with environmental sustainability goals. This trend indicates a movement towards addressing not just pollutant removal, but also harnessing by-products (such as methane) in the process, thereby contributing to a circular economy approach.

The frequency data highlights some core topics that have sustained attention over time. “*Bioaugmentation*” (249 occurrences) and “*degradation*” (144 occurrences) are particularly prominent, emphasizing the consistent research interest in enhancing microbial communities to break down pollutants effectively. The significant frequency of “*biodegradation*” (102 occurrences) and “*methane production*” (125 occurrences) further underscores a dual focus on both pollutant degradation and energy/resource recovery.

Figure 3b provides a comprehensive overview of research trends in environmental biotechnology over nearly two decades. It highlights the diverse journals through which knowledge has been disseminated and illustrates the geographical distribution of research efforts. The figure effectively portrays the global landscape of contributions, emphasizing the interconnectedness of leading countries, key research topics, and significant scientific publications in the field. Using a Sankey diagram, it highlights how major research areas such as “bioaugmentation,” “anaerobic digestion,” “bioremediation,” and “microbial community” studies are distributed across various countries and published in specific journals. China and the USA are identified as the most significant contributors, with substantial research outputs across multiple topics, followed by other active countries like India, Korea, and Italy. The diagram reveals that *Bioresource Technology*, *Environmental Science & Technology*, and *Water Research* are among the top journals publishing these studies, with *Bioresource Technology* standing out as a primary publication venue across diverse topics. The interconnected flows in the diagram emphasize how different countries focus on similar research areas and target common journals, creating a cohesive international research network. This visualization effectively captures the interdisciplinary and collaborative nature of environmental biotechnology research, underscoring the global commitment to advancing sustainable biotechnological solutions.

**Figure 3.**
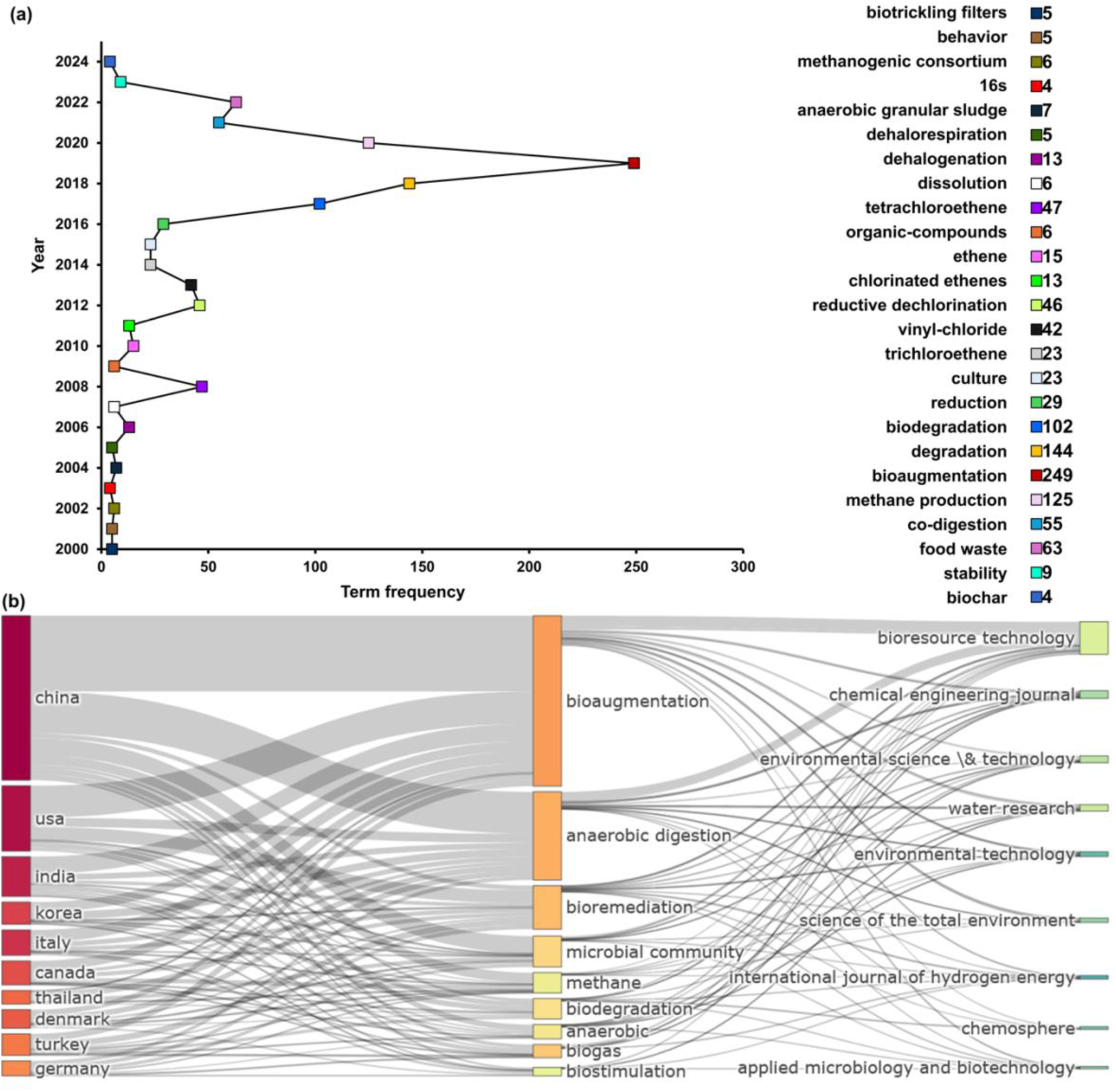
Key word analysis, journals and geographic distribution: (a) frequency of keywords from the years 2000 to 2024. The right side displays the ranking of each term by frequency, reflecting their prominence in the literature; (b) Sankey diagram illustrating the prominent countries, popular keywords, and frequently cited journals in the field. Both visualizations were prepared using the R package *bibliometrix*.

**Figure 4.**
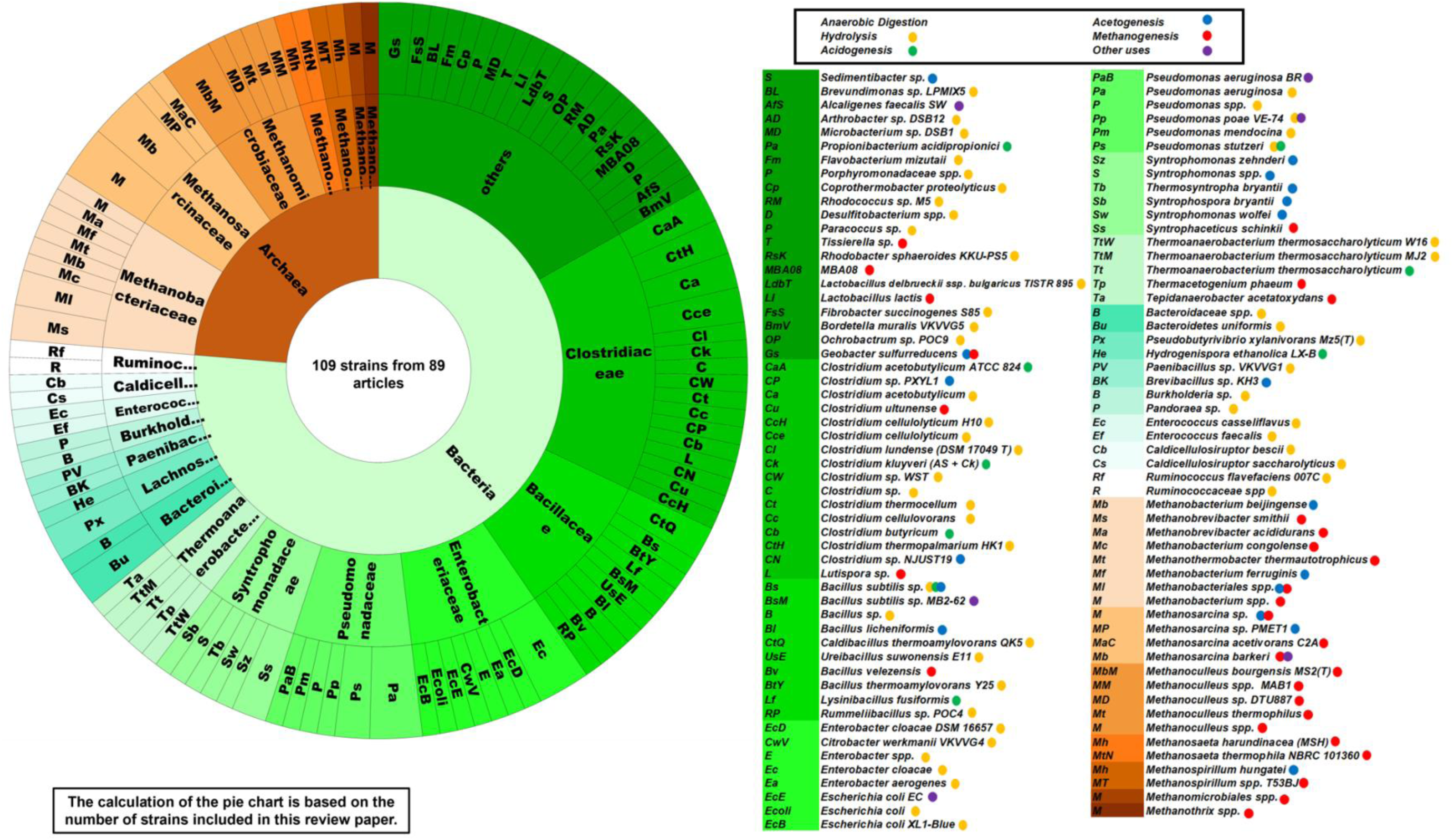
Pie chart indicating the diversity and frequency of microbial strains used for bioaugmentation in anaerobic digestion systems. In the figure and throughout the study, the affiliations *Methanothrix* and *Methanosaeta* have both been used. It must be noted that more recent works use *Methanothrix* instead of *Methanosaeta*, as *Methanosaetaceae* were renamed to *Methanotrichaceae*.

### 3.2 Differentiation from existing review articles

While numerous review articles have been published on the topic of anaerobic digestion and bioaugmentation, this systematic review distinguishes itself by addressing critical gaps in the current literature. Many existing reviews provide comprehensive overviews of anaerobic digestion processes and the general application of bioaugmentation. However, they often lack a detailed analysis of the different control conditions under which bioaugmentation is implemented, and they do not adequately differentiate the specific roles of various microorganisms at each step of the anaerobic digestion process.

In anaerobic digestion, the microbial consortia involved are diverse and perform distinct functions at various stages, such as hydrolysis, acidogenesis, acetogenesis, and methanogenesis. While some reviews touch on the role of microorganisms in enhancing specific steps of bioprocesses, there remains a lack of comprehensive discussion on their effectiveness across various operational conditions. This review fills this gap by systematically identifying key microorganisms involved in different bioprocess steps and providing insights into how their effectiveness can vary under different substrates, environmental settings, and reactor configurations. By offering a more nuanced understanding of these factors, this work contributes to refining bioaugmentation strategies for diverse operational contexts. This review aims to build upon previous works by systematically analyzing the impact of various control conditions, such as pH, temperature, and inoculum sources, on the effectiveness of bioaugmentation. While there have been studies addressing aspects of bioaugmentation, our approach differentiates the microbial roles and optimizes conditions for each stage of anaerobic digestion in a comprehensive manner. By focussing on taxonomy differentiating the microbial roles and optimizing conditions for each stage of anaerobic digestion, this review provides a more targeted approach to improving the efficiency and effectiveness of bioaugmentation in anaerobic digester plants.

### 3.3 Controls used in bioaugmentation experiments

In this section the focus lays on the various measures implemented to control the growth and activity of added microbes in the context of bioaugmentation experiments. These controls are crucial for ensuring reliable and accurate results while studying the potential of bioaugmentation for enhancing biological processes. The initial aim of this section was to highlight, which controls were used to demonstrate that bioaugmentation resulted in increased biogas levels. However, it turned out that the diverse set-ups of different experiments make the categorization difficult, especially since not all experiments aimed to produce methane. Originally, it was planned to divide all experiments into cases with (1) no control, (2) control without adding additional bioaugmentation strains, (3) controls, where autoclaved strains were added, (4) positive controls with strains, which are known to give positive results. However, it turned out that the applied search sometimes resulted also in some extraordinary approaches, which were dealing with rather unusual scenarios (e. g. the screening for new strains for bioaugmentation or MFCs. For these (5) extraordinary approaches a fifth type of control has been defined, which is further referred to as “other”. All found articles have been assigned to these five groups of controls (Table 1). In several articles, insufficient control experiments where observed. In several articles, no control was used. Usually, there is only one type of control applied (control group 2, 3 or 4). On few occasions, two types were applied. In none of the studies, all three types of controls were applied.

**Table 1:**
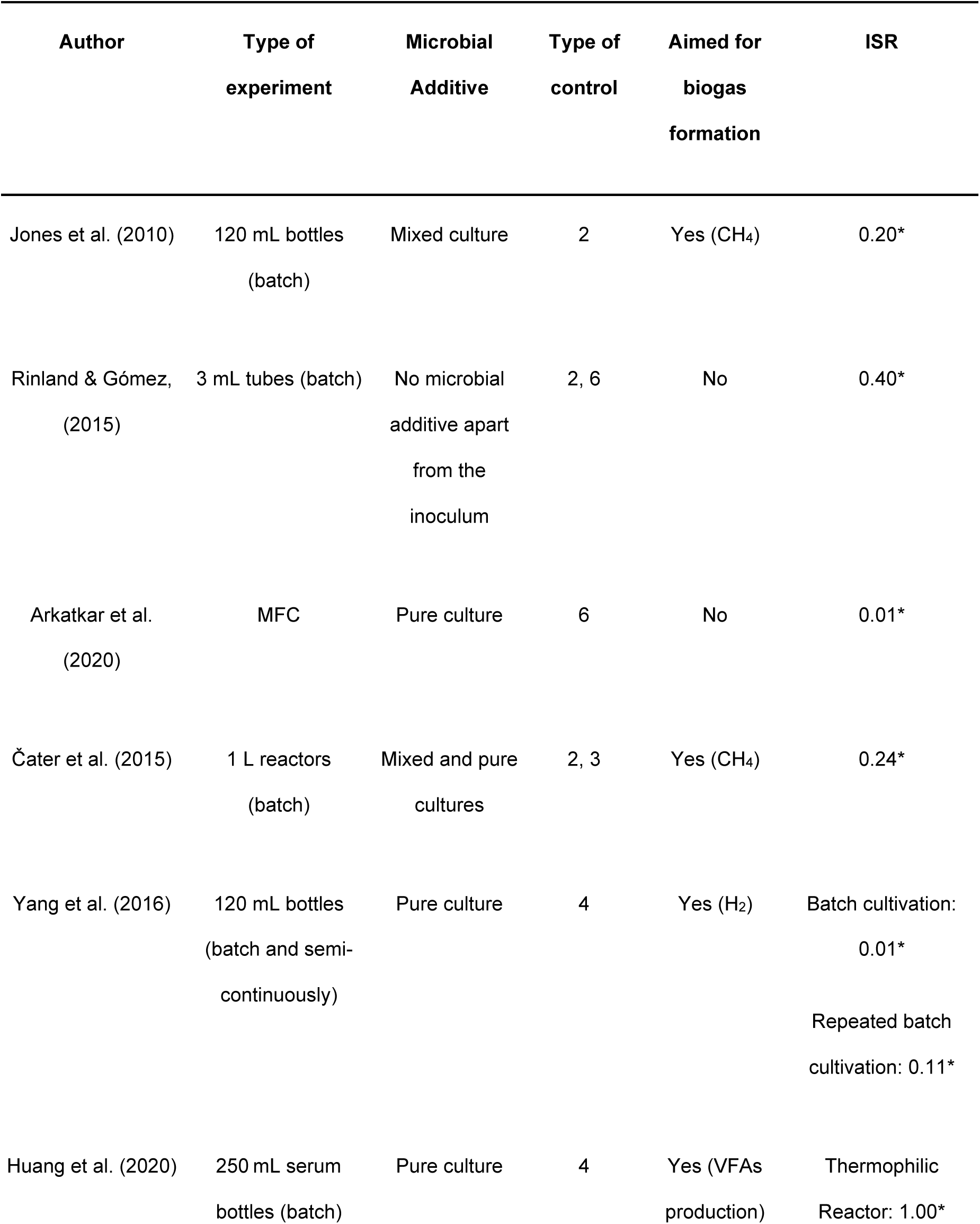
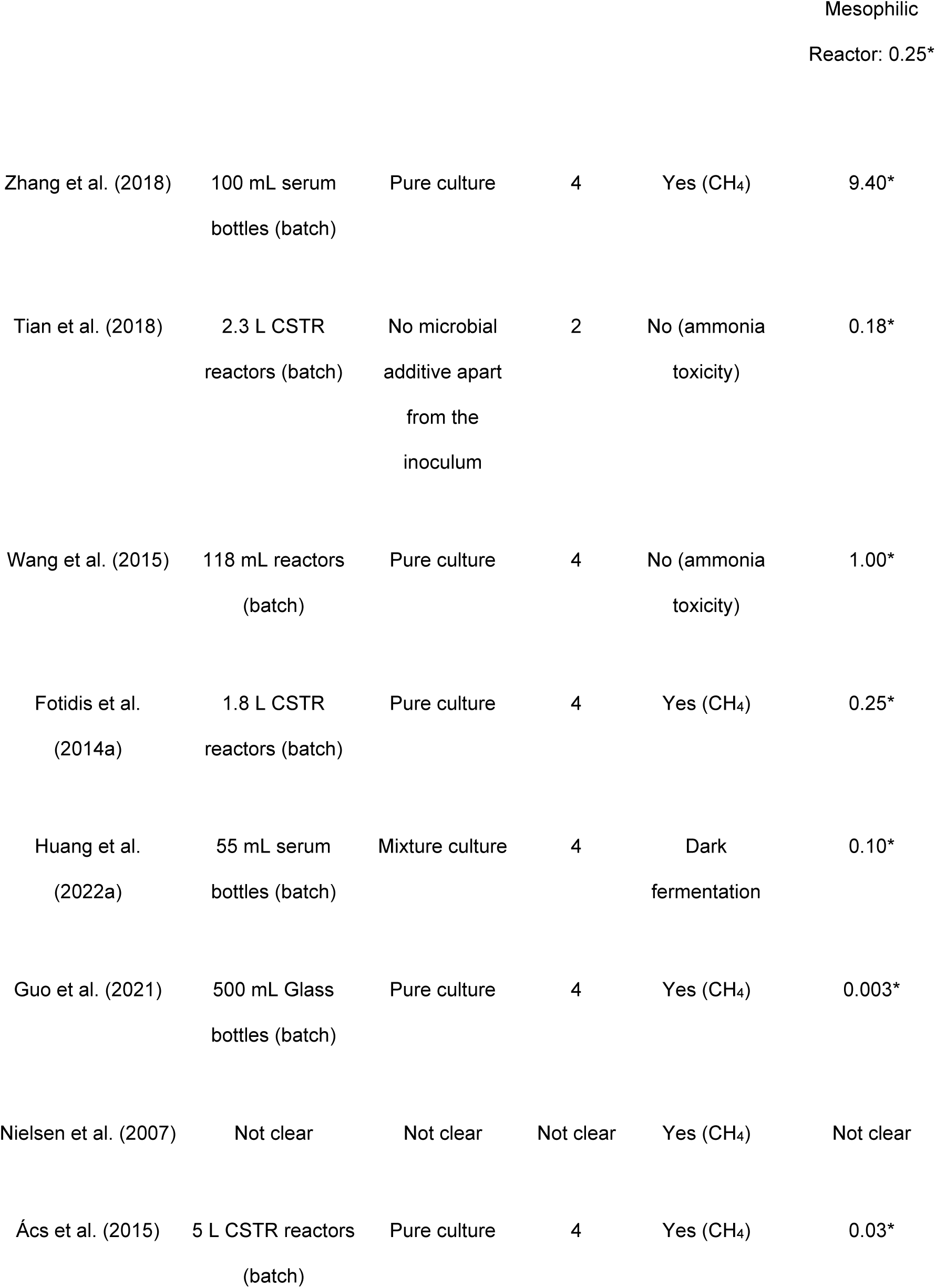
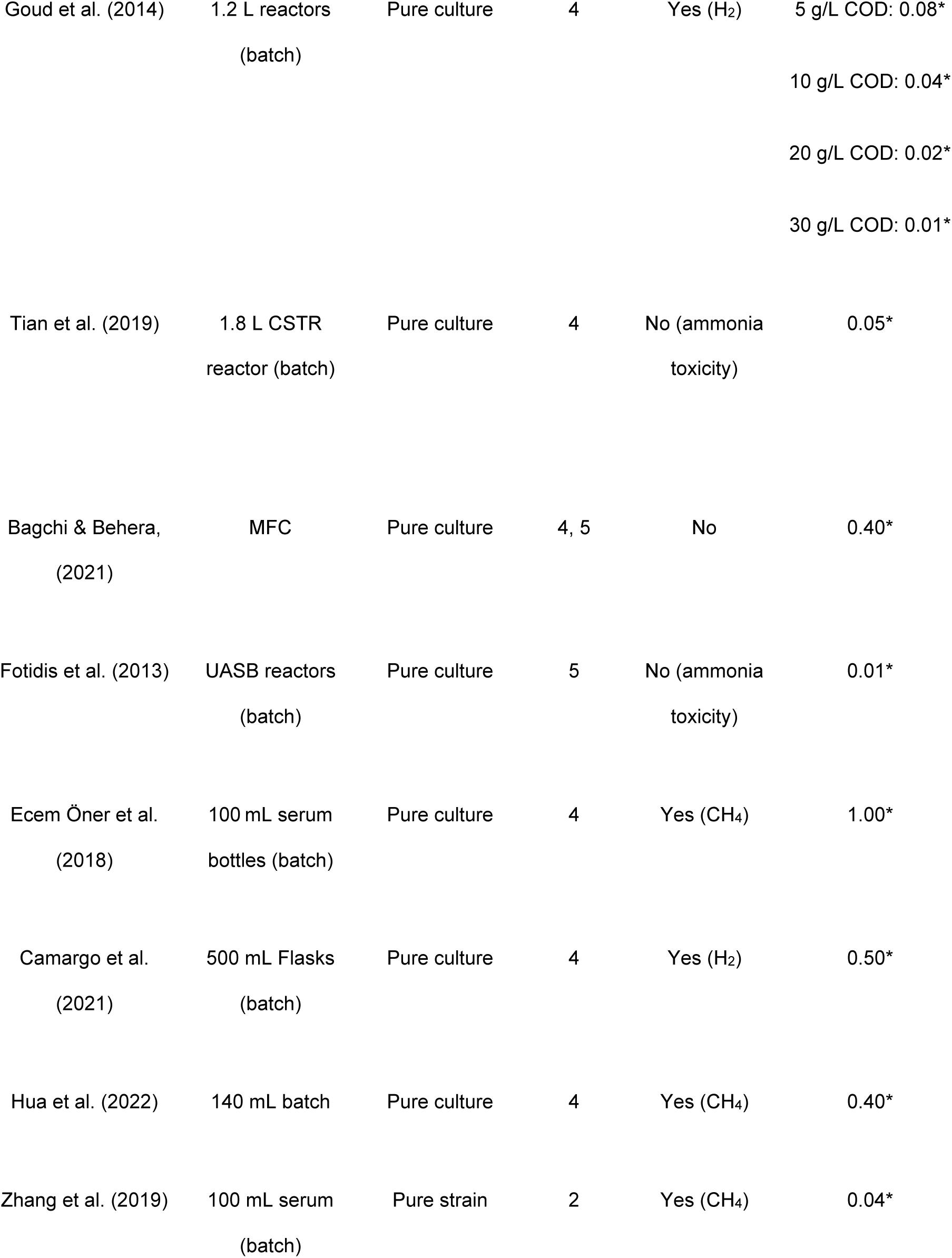
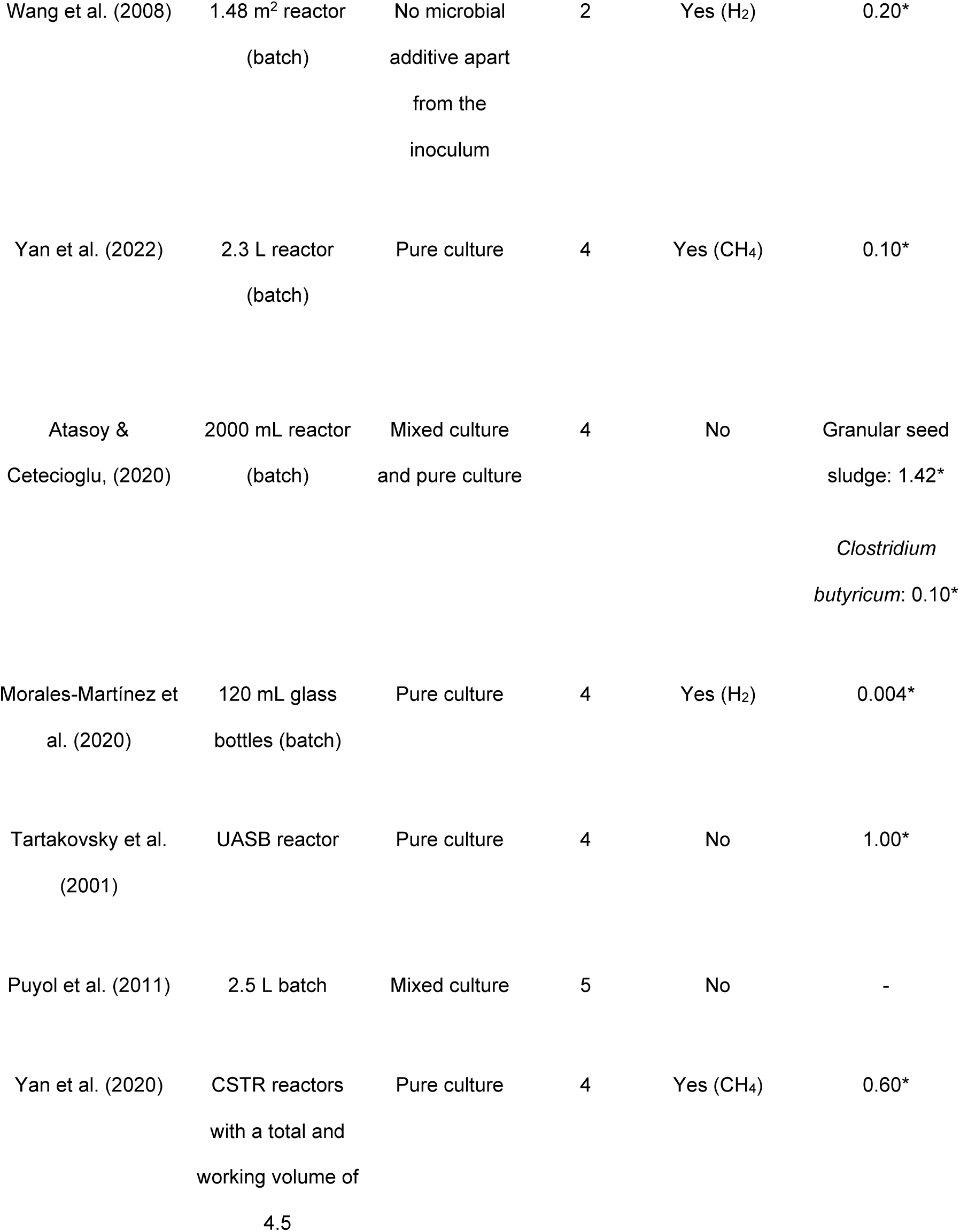
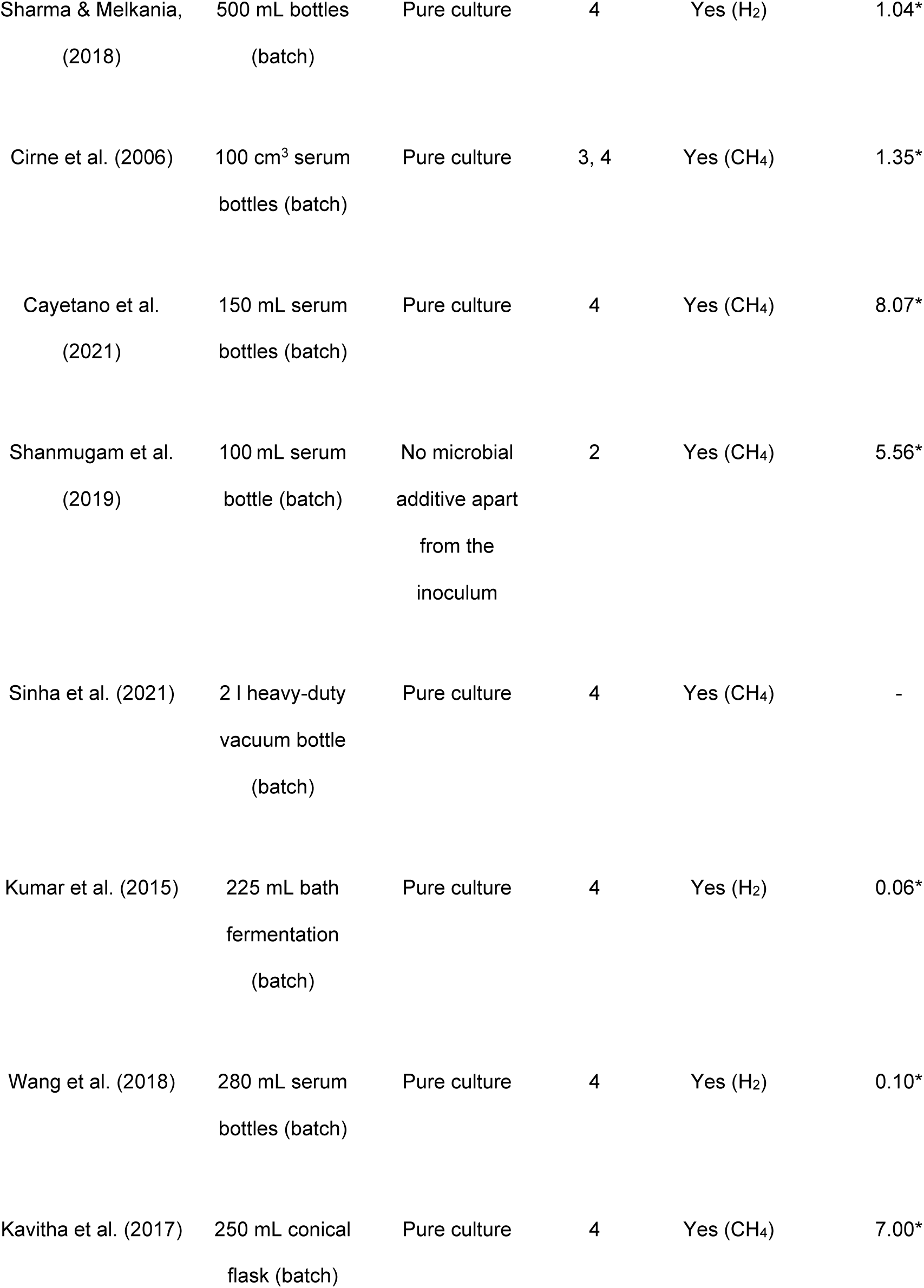
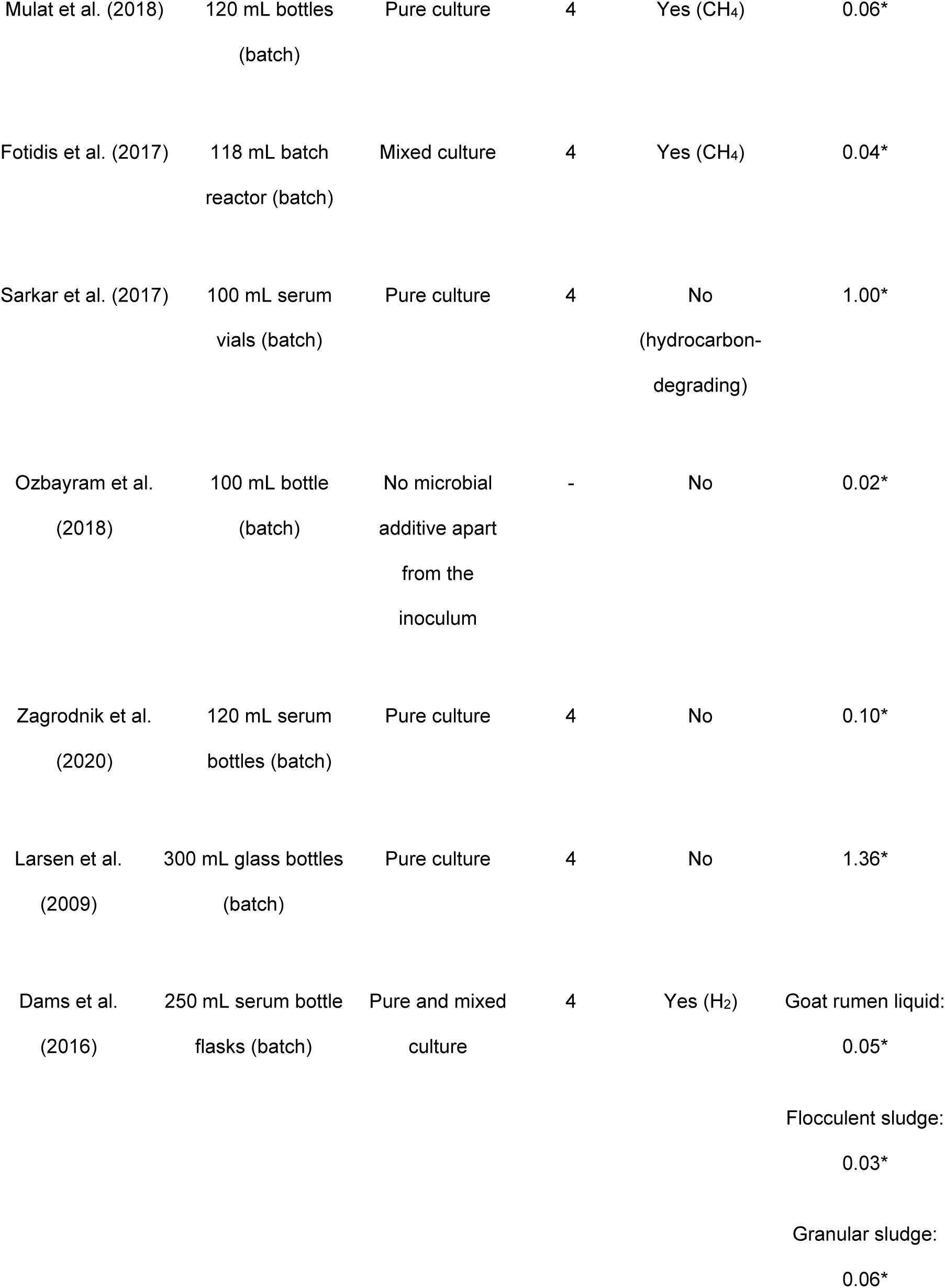
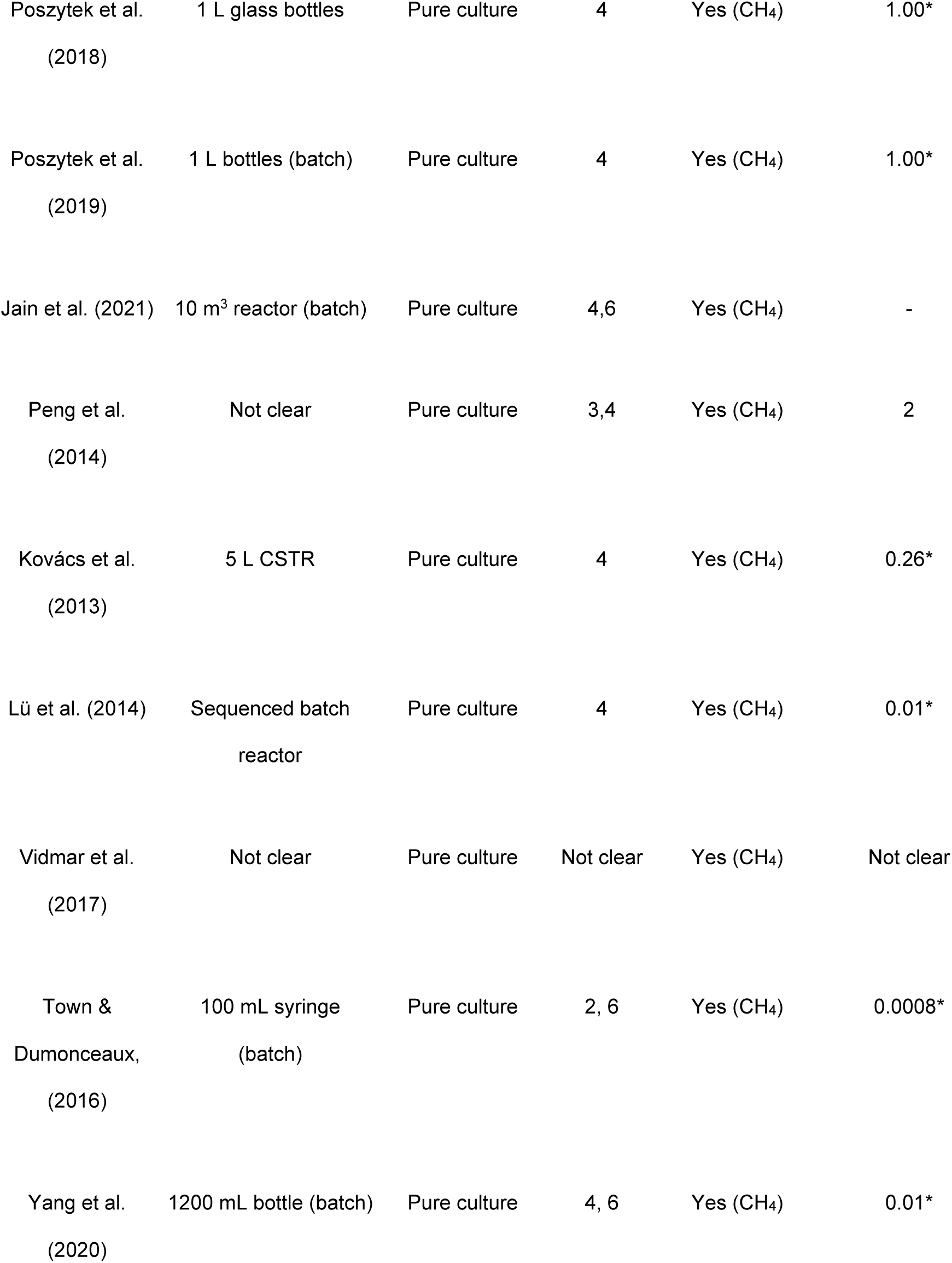
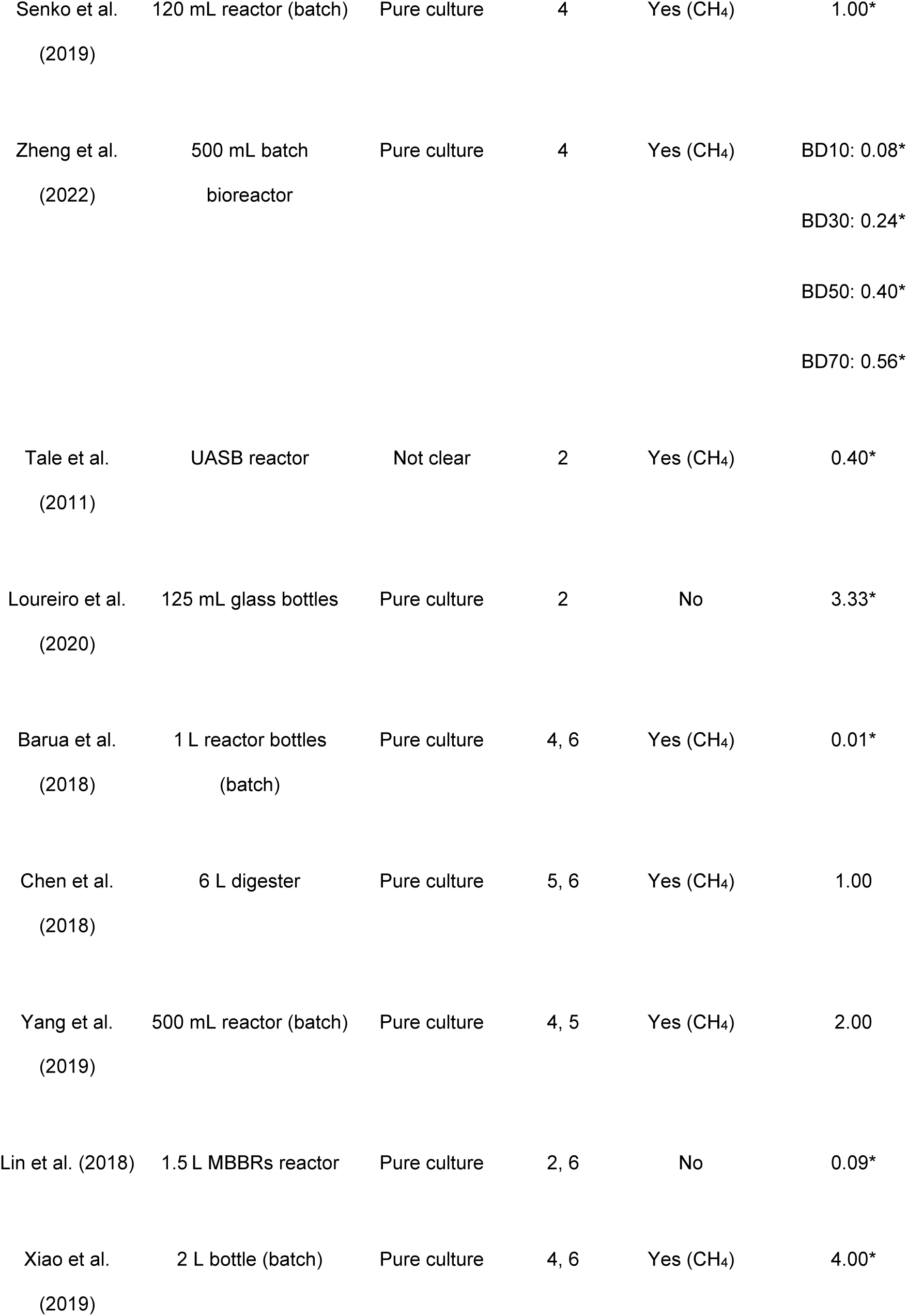
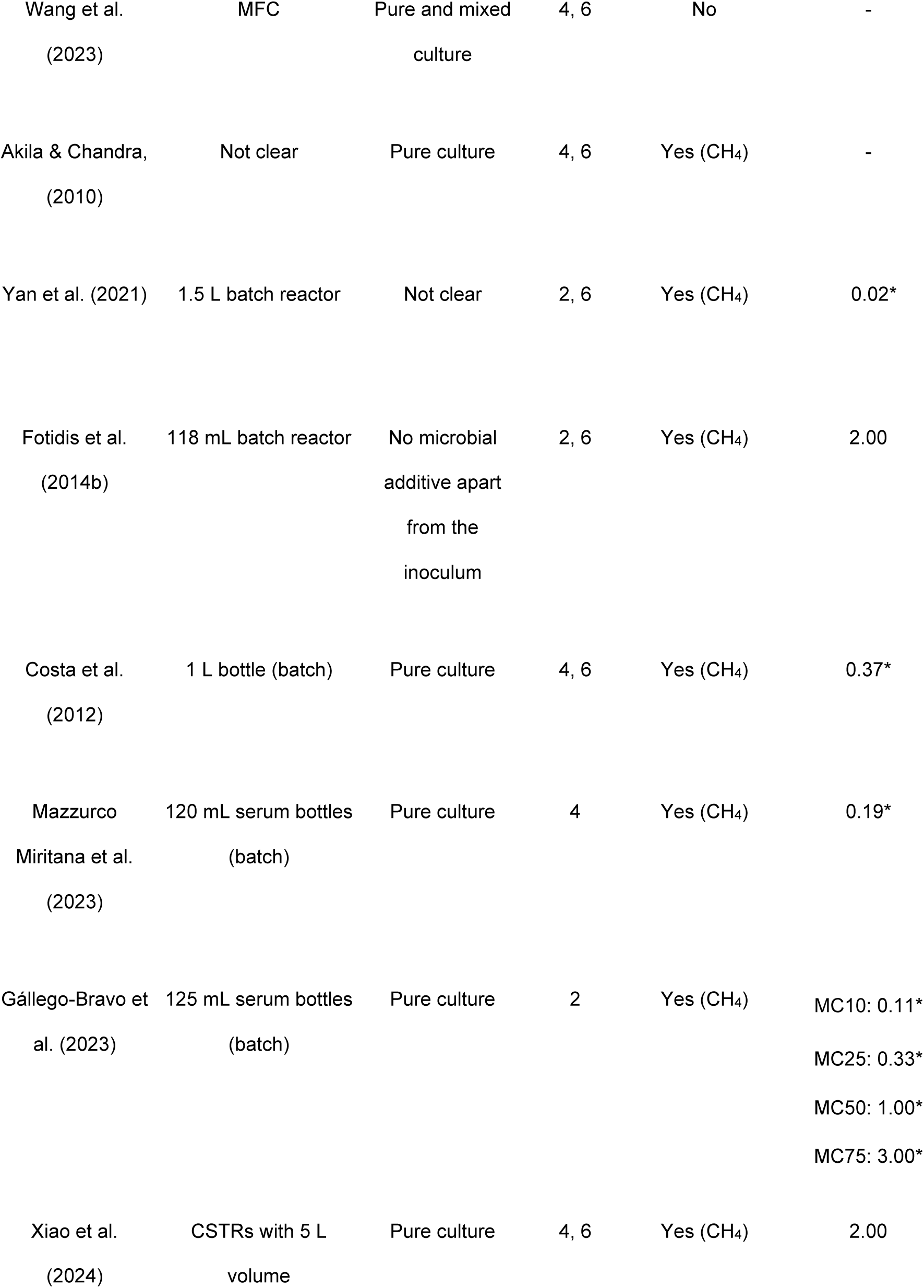
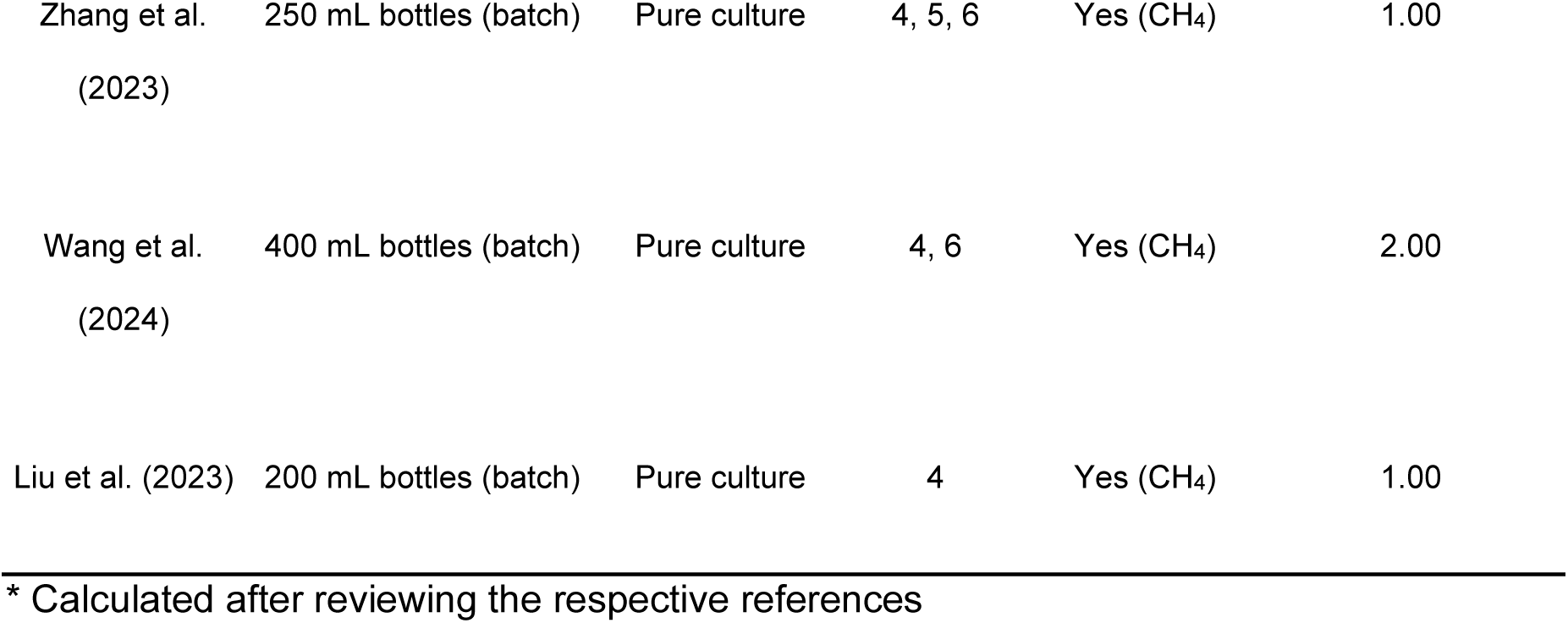
Types of controls, which have been used in publications on bioaugmentation: controls have been assigned as (1) no control, (2) control without adding additional bioaugmenting strains, (3) controls, were autoclaved strains were added, (4) positive controls with strains, which are known to give positive results, (5) negative controls with strains, which are known to give ineffective results other, (6) (approaches with rather unusual scenarios). Not all 89 articles are shown, since full text where not available for all articles.

Comparing the “types of experiment” in table 1, it becomes clear that the Biochemical Methane Potential (BMP) assay is common choice. This assay is a valuable method for determining the ultimate biodegradability and methane conversion yield of organic substrates (Angelidaki et al., 2009). A critical parameter in the BMP assay is the inoculum-to-substrate ratio (ISR), which significantly influences the efficiency of anaerobic degradation, the relevance of the degradation test to full-scale digesters and the accuracy of the assay. Research has shown that a higher ISR can improve the ultimate practical methane yield. For example, a batch digestion test on microalgae found that an ISR of 2, compared to 1 and 0.33, resulted in the highest methane productivity, ranging from 188 to 395 mL CH₄/g VS added across different microalgae types (Alzate et al., 2012). The digestion of sunflower oil cake (SuOC) at an ISR of 3, compared to lower ratios, produced the highest methane yield (Raposo et al., 2009). However, at lower ISRs, while the maximum specific methane production rate was higher, the overall methane yield was lower, as observed in BMP tests of maize at various ISRs (Raposo et al., 2009). This lower yield at low ISRs was linked to the accumulation of longer-chain acids within the system, which could inhibit methanogens, particularly due to the acetate produced during digestion at high substrate concentrations (Maya-Altamira et al., 2008). Increasing the ISR, which involves diluting the substrate, can help enhance practical methane yield. Most studies have focused on the impact of ISR on methane yield for single substrates, with limited documentation on its effects in co-digestion scenarios. Additionally, the source of inoculum is crucial, especially when dealing with complex substrate mixtures, due to the diverse microbial consortia involved. Calculating the ISR is important because it directly impacts the methane production efficiency and overall yield, which is illustrated in table 1. Different substrates produce varying methane outputs, which can be effectively assessed by considering the ISR in the BMP assay. It is hypothesized by the authors a low ISR, or usage of an inoculum source that is unsuitable for the substrate of choice can lead to false-positives on the effect of bioaugmentation.

The ISR values presented in Table 1 show significant variability across different studies, reflecting diverse experimental setups and microbial additives. In this review only 34% of the studies meet the minimum required ISR of >1, and only 15% of the studies meet the minimum desired ISR of >2. This threshold is based on Holliger et al. (2016). Among the entries, the highest ISR is reported by Zhang et al. (2018), with a value of 9.4, achieved using a pure culture in 100 mL serum bottles aimed at biogas production. This indicates that the selected microbial strains and experimental conditions can greatly influence ISR outcomes. In contrast, studies such as Arkatkar et al. (2020) report much lower ISR values, such as 0.01, due to the use of a pure culture in a microbial fuel cell (MFC) setup. The discrepancies in ISR values across authors can be attributed to differences in the type of cultures used (mixed vs. pure), the experimental design (batch vs. continuous systems), and the specific aims of the research, such as methane or hydrogen production. Therefore, while some ISRs suggest strong potential for biogas formation, others indicate challenges that may require further optimization or different microbial approaches. Overall, identifying the most effective ISR for biogas production depends on the specific research context and microbial strains utilized.

### 3.4 Manipulation of hydrolysis

The present work tries to distinguish applied microbes according to the different phases of anaerobic digestion (hydrolysis, acidogenesis, acetogenesis and methanogenesis). This separation is not always feasible, as there are overlaps. For example, some hydrolytic bacteria yield organic acids, which results in an overlap between hydrolysis and acidogenesis. This simultaneous involvement blurs the boundaries between the hydrolysis and acidogenesis phases. Similar challenges arise in other phases, such as acetogenesis, where certain microorganisms may contribute to both acidogenesis and acetate production, creating further complexities in differentiation. The difficulty of dividing found articles into the various phases of anaerobic digestion is also made clear in a work by Zhang et al. (2019). Zhang et al. (2019) used the hydrolytic *Thermoanaerobacterium thermosaccharolyticum W16* mixed with undefined methanogenic granular sludge. It did not only improve hydrolysis, but also syntrophic relations, which are rather related to the later phases of anaerobic digestion. Nevertheless, the authors of the present study tried to distinguish the phases as clearly as possible, starting with hydrolysis. Hydrolysis contemplates the first stage of anaerobic digestion. During hydrolysis, complex organic compounds (e.g., carbohydrates, proteins, and lipids) will be transformed into simpler molecules, like sugars, long chains of fatty acids and amino acids due to the enzymatic attack made by different types of anaerobic microorganisms. During this stage, various obstacles may arise that limit the AD process, as well as the performance and adequate production of biogas.

The use of bioaugmentation within hydrolysis has been studied for various purposes with the overall goal of improving the efficiency and overall stability of the AD process. The systematic literature search performed in the present work resulted in 33 articles, which were predominantly focused on the inoculation of hydrolytic microorganisms for bioaugmentation within the first stage of AD. Many of them are focussed on the improved degradation of fibre-rich material. For example, two of them highlighted the use of bioaugmentation to increase the methane yield from cattle manure and brewery spent grain. Hydrolytic organisms are promising here, as the mentioned substrates contain lignocellulosic biomass, which usually degrades very slowly and, therefore, the hydrolysis takes longer to complete (Nielsen et al., 2007; Čater et al., 2015). In a similar approach, improved lignocellulose degradation has been shown with *Citrobacter werkmanii VKVVG4, Bordetella muralis VKVVG5* and *Paenibacillus sp. VKVVG1,* but for water hyacinth (Barua et al., 2018). Another promising approach was presented by Peng et al. (2014), who was able to enhance wheat straw hydrolysis and to improve the biochemical methane potential (BMP) from wheat straw due to the application of the cellulolytic anaerobic bacterium *Clostridium cellulolyticum*. The outcomes showed BMPs of 342.5 ml g^-1^ VS and 326.3 ml g^-1^ VS, representing a 13.0% and 7.6% increase, respectively, compared to the BMP without bioaugmentation, which was 303.3 ml g^-1^ VS. Similar to Peng et al., Ecem Öner et al. (2018) worked on the degradation on wheat straw too. They used *Clostridium thermocellum* to enhance methane yield from lignocellulosic biomass by up to 39%. Ozbayram et al. (2018), also worked with wheat straw as a substrate, enriching methanogenic communities from cow and goat rumen fluid and a biogas reactor. The dominant strains in the enriched cultures were *Bacteroidaceae spp*. (rumen) and *Porphyromonadaceae spp.* (reactor), with an increased abundance of *Ruminococcaceae spp.* (Firmicutes). Similarly, Sinha et al. (2021) employed the cellulolytic strains *Microbacterium sp. DSB1* and *Arthrobacter sp. DSB12* for lignocellulose degradation of *Lantana camara*, achieving enhanced biogas production with methane yields of 57% and 60%, respectively. Based on the afore mentioned articles it stands out that especially bacteria from the phylum Firmicutes are used abundantly to improve the degradation of lignocellulose. In this regard, another study by Shanmugam et al. (2019) can be highlighted, which focused on the strain *Clostridium sp. WST.* After isolating it from mangrove sediments, this strain improved the degradation of lignocellulosic biomass degradation due to bioaugmentation in anaerobic digestion experiments.

Another article, where cellulolytic bacteria (*Bacillus sp.*) were applied too, has been presented by Kavitha et al. (2017). The work stands out as the experiment was not just focussed on improved hydrolysis from substrates within a given methanogenic digester. Instead, the substrate was pretreated before it was entered into the respective reactor. In this specific case, Kavitha et al. (2017) investigated the impact of bacterial-based biological pretreatment on the liquefaction of *Chlorella vulgaris* microalgae before anaerobic biodegradation. The results show that pretreatment with cellulose-secreting bacteria increase the biomass stress index by 18% compared to the control group. The biomass stress index measures the physical and chemical stress experienced by the biomass during the pretreatment process, indicating the extent to which the biomass is broken down and made more amenable to subsequent processes. In this study, pretreatment significantly enhanced the biomass stress index, demonstrating its effectiveness in preparing Chlorella vulgaris for anaerobic biodegradation. Similar to Kavitha et al., Mulat et al. (2018) and Vidmar et al. (2017) tried to improve biomass pretreatment too. For this, Mulat et al. combined steam explosion (SE) and bioaugmentation, which significantly enhanced methane yield from birch. Compared to the untreated control, the yield increased up to 140%. Bioaugmentation with *Caldicellulosiruptor bescii* at lower dosages (2% and 5% of inoculum volume) showed the best methane improvement on day 50. Additionally, the microbial community analysis indicated an increase in abundance of key bacterial and archaeal communities, including the hydrolytic bacterium *Caldicoprobacter*, syntrophic acetate oxidising bacteria, and hydrogenotrophic *Methanothermobacter*, contributing to the enhanced methane production. Vidmar et al. (2017) utilized the anaerobic bacterium *Pseudobutyrivibrio xylanivorans Mz5^T^* to pretreat biomass under mesophilic and thermophilic conditions. This pretreatment targeted the hydrolysis of resistant cell wall components, such as cellulose and hemicellulose. Additionally, Costa et al. (2012) used the strains *Clostridium cellulolyticum*, *Caldicellulosiruptor saccharolyticum*, and *Clostridium thermocellum* for biological co-treatment as a pretreatment for enhancing the degradation of poultry litter.

To investigate the potential of bioaugmentation by anaerobic hydrolytic bacteria, Čater et al. (2015) conducted a study that included a BMP assay. Active microorganisms from a full-scale upflow anaerobic sludge blanket reactor treating brewery wastewater, along with brewery spent grain as a representative lignocellulosic substrate, were used pure and mixed cultures of *Ruminococcus flavefaciens 007C*, *Pseudobutyrivibrio xylanivorans Mz5^T^*, *Fibrobacter succinogenes S85*, and *Clostridium cellulovorans* were employed to enhance lignocellulose degradation and increase biogas production. *P. xylanivorans Mz5^T^* exhibited the highest methane production increase (+17.8%), followed by specific co-cultures that also demonstrated improvements. In regard to hydrolysis and compared to the upper examples on bioaugmentation, the work from Čater et al stands out as it highlights the possibility to combine more than one microbe on a combined approach. Fingerprinting techniques revealed significant changes in the microbial community structure, highlighting the impact of the experimental conditions on microbial dynamics. Nevertheless, Čater et al. are not the only authors, which work with multi-species bioaugmentation. Similarly to Čater et al., Senko et al. (2019) employed *Clostridium acetobutylicum*, *Pseudomonas sp*., and *Enterococcus faecalis* not only for lignocellulosic waste but also for substrates containing antibiotics and pesticides. With the combination of strains, biogas production significantly increased, demonstrating the effectiveness of immobilized anaerobic sludge cells in enhancing methanogenesis.

Apart from tackling lignocellulose degradation, there are other substrates which are difficult to degrade, such as biodiesel, which has been addressed by Sarkar et al. (2017). For this, they performed bioaugmentation with *Enterobacter*, *Pandoraea*, and *Burkholderia* strains in order to effectively biodegrade hydrocarbons. Also working with diesel, Loureiro et al. (2020) has shown the suitability of *Pseudomonas aeruginosa*, *Pseudomonas stutzeri*, and *Pseudomonas mendocina*. It is further possible to lower negative impacts from contaminations, e.g. with heavy metals. In a study by Guo et al. (2021), *Paracoccus* sp. was utilized in bioaugmentation experiments with plant residues rich in heavy metals, significantly enhancing degradation efficiency and boosting biogas and methane production. In the studies by Puyol et al. (2011) and Tartakovsky et al. (2001), bioaugmentation was tested to improve the degradation of toxic compounds in anaerobic systems. Puyol et al. used *Desulfitobacterium* strains to bioaugment the degradation of 2,4,6-trichlorophenol (246TCP), a chlorinated pesticide, but found no significant improvement in its anaerobic biodegradation. Similarly, Mo et al. (2011) bioaugmented reactors with the aerobic biphenyl degrader *Rhodococcus sp. M5* for the degradation of Aroclor 1242, a PCB mixture, but observed no enhanced performance, with similar degradation rates in both bioaugmented and non-bioaugmented reactors.

So far, the majority of the studies were focussed on the co-digestion of biomass in classical co-digesters. However, and apart from this, hydrolytic strains hold further the potential to improve other processes such as wastewater treatment. In this regard, a promising approach of bioaugmentation was presented for the anaerobic digestion of waste-activated sludge, where the hydrolytic bacteria, *Bacteroidetes uniformis* and *Clostridium sp.* were introduced at different dosages. The bioaugmentation resulted in a remarkable enhancement in methane conversion from waste-activated sludge. In another study on wastewater treatment conducted by Poszytek et al. (2018, 2019), the use of *Rummeliibacillus sp. POC4*, *Ochrobactrum sp. POC9*, and *Brevundimonas sp*. *LPMIX5* was investigated for enhancing hydrolysis and biogas production. The results demonstrated a significant increase of 22% and 28% in hydrolysis and biogas production, respectively, when *Rummeliibacillus sp. POC4* and *Ochrobactrum sp. POC9* were used with sewage sludge. The most significant methane yield of 298.1 mL CH_4_/g COD, with an impressive 85.2% COD conversion efficiency, was achieved when *Bacteroidetes uniformis* and *Clostridium sp*. were added at 100 and 900 CFU/mL, respectively (Cayetano et al., 2021). In regard to wastewater treatment, another interesting article can be found by Li et al. (2010). Li et al. applied bioaugmentation to enhance the cellulose degradation capacity of an upflow anaerobic sludge blanket (UASB) reactor utilized for treating swine wastewater, a composite microbial consortium was introduced as a bioaugmentation strategy. The objective was to bolster the reactor’s efficiency in breaking down cellulose, a complex organic compound found in the wastewater. The results indicated that the microbial community structure changed significantly, with the inoculated bacteria *Flavobacterium mizutaii* and *Pseudomonas* as the dominant strains in the reactor. Due to bioaugmentation, bacterial populations such as *Clostridia*, *Acidobacteria*, and *Nitrospira* were present in the bio-augmented system. Dominant groups like *Chloroflexi* and *Acidobacteria*, and several groups of *Bacteroidetes*, *Proteobacteria*, and *Firmicutes* either had much lower density or disappeared in the incubated bacterial system. By incorporating this specialised consortium of microorganisms into the reactor’s microbial community, it was intended to improve the overall degradation performance and optimise the treatment process (Li et al., 2010). In regard to wastewater treatment it needs to be highlighted that there are cases, which can’t be attributed to any of the four phases. In this regard, a study by Lin et al. (2018) can be mentioned, which used *Pseudomonas aeruginosa* for denitrification treatment processes.

A very interesting approach was shown by Lü et al. (2014). They focused on the degradation of sludge digestate through thermophilic anaerobic digestion with the addition of thermophilic, proteolytic *Coprothermobacter proteolyticus* and/or methanogenic granular sludge. The application of a proteolytic microbe is interesting, as this shows an alternative approach to the cellulolytic approaches further above. In the study by Lü et al., the sludge digestate stabilized by mesophilic anaerobic digestion was further degraded through thermophilic anaerobic digestion using 0–10% (v/v) of thermophilic, proteolytic *Coprothermobacter proteolyticus*, and/or methanogenic granular sludge. This optimization step, conducted prior to the bioaugmentation, demonstrated that the temperature shift to thermophilic conditions promoted abiotic solubilization of proteins and reactivated fermentative bacteria and methanogens indigenous to the sludge digestate, resulting in a final methane yield of 6.25 mmol-CH_4_/g-volatile suspended solid (VSS) digestate. The inclusion of *Coprothermobacter proteolyticus* accelerated hydrolysis and fermentation during the early stages of thermophilic anaerobic digestion and stimulated methane production through syntrophic cooperation with methanogenic granular sludge, achieving a final methane yield of 7 mmol-CH_4_/g-VSS digestate with significant protein and polysaccharide degradation.

Next to lignocellulose and refractory proteins, the successful degradation of lipids is a challenge too. In this regard, a promising approach of bioaugmentation was presented in a study conducted by Cirne et al. (2006). They used the lipolytic bacterial strain *Clostridium lundense (DSM 17049T), which* was found to enhance methane production during anaerobic digestion of lipid-rich waste. The research demonstrated a higher methane production rate of 27.7 cm^3^ CH_4 (STP)_ g^-1^ VS _added_ day^-1^ (VS, volatile solids) under methanogenic conditions. Moreover, the bioaugmentation strategy significantly improved the hydrolysis of the lipid fraction, evident from the highest initial oleate concentration of 99% in the substrate. Xiao et al. (2024) investigated lipid-rich substrates as well. In their particular case, they investigated the effects of microbial bioaugmentation on methane production in thermophilic anaerobic digestion (TAD) from food waste. Similar to Čater et al. (2015), they performed bioaugmentation with an inoculum consisting of more than one strain. They used a combined inoculant of *Clostridium thermopalmarium HK1* and *Caldibacillus thermoamylovorans QK5*, which improved cumulative methane production by 24.77% compared to the control. Interestingly, the combined inoculant enriched carbohydrate- and protein-degrading bacteria, boosting carbohydrate metabolism, amino acid metabolism, and methane metabolism. This strategy specifically enhanced the methanogenesis step by promoting the tricarboxylic acid cycle (TCA cycle) and the conversion of CO_2_ to methane. So far, all approaches focused on the improved hydrolysis of various substrates. However, and especially in the case of separate hydrolysis stages, hydrogen production is another important objective. It could be assigned to both, hydrolysis and acidogenesis. An example for this is the work by Mazzurco Miritana et al. (2023), who tested bioaugmentation with a fermenting bacteria pool and anaerobic fungi (*Orpynomyces sp. and Neocallimastix sp.*) to enhance both, the “fermentative and hydrolytic phases” during anaerobic digestion of shrimp processing waste. Another example, where both phases are addresses, has been published by Zhang et al. (2019). They explored bioaugmentation with *Thermoanaerobacterium thermosaccharolyticum W16* to enhance thermophilic hydrogen production from corn stover hydrolysate. The addition of a small amount of strain *W16* (5% of total microbes) resulted in increased hydrogen yields across different seed sludge types. This bioaugmentation process also influenced the composition of soluble metabolites, favouring acetate production and reducing butyrate and ethanol accumulation in specific situations. Microbial community analysis revealed the dominance of *Thermoanaerobacterium spp.* and *Clostridium spp.* The abundance of Thermoanaerobacterium indicates that this genus was enriched successfully in the respective reactor due to bioaugmentation. It is interesting that apart from *Thermoanaerobacterium* the genus *Clostridium* was increased too, which suggests a vital role for this genus in thermophilic hydrogen generation. For some substrates, such as rotten corn stover and sludge from anaerobic digestion, the bioaugmentation significantly increased the relative abundance of strain *T. thermosaccharolyticum W16* in the microbial community, likely contributing to the enhanced hydrogen production as well. In a similar study Huang et al. (2022a) investigated the use of *T. thermosaccharolyticum MJ2* and biochar to enhance thermophilic hydrogen production from sugarcane bagasse. The study found that bioaugmentation with *MJ2* significantly increased hydrogen production by 95.31%, and the addition of biochar further enhanced this by an impressive 158.10%. In addition, Camargo et al. (2021) used a strain closely related to *Enterococcus casseliflavus* for hydrogen production. The strain was isolated from citrus by-products and demonstrated significant hydrogen production from xylose. Additionally, In a study by Kumar et al. (2015), bioaugmentation with *Escherichia coli XL1-Blue* and *Enterobacter cloacae DSM 16657* significantly improved hydrogen production from beverage industrial wastewater. The addition of facultative anaerobic bacteria, combined with nutrients such as yeast extract and tryptone, led to a remarkable increase in hydrogen production, especially when both the bacteria and nutrients were used together. As mentioned already further above, it can be promising to use more than one strain in a combined approach. This is also possible in hydrogen production. In this regards, Laocharoen et al. (2015) investigated bioaugmentation for hydrogen production by adding *Rhodobacter sphaeroides KKU-PS5* and *Lactobacillus delbrueckii ssp. bulgaricus TISTR 895* into anaerobic digesters. While the co-cultivation faced challenges due to differences in metabolic types, this study highlighted the potential of combining these strains to improve hydrogen production through bioaugmentation. In a similar approach, Sharma & Melkania, (2018) evaluated the effect of bioaugmentation with three bacterial species (*Escherichia coli, Bacillus subtilis*, and *Enterobacter aerogenes*) on hydrogen production from the organic fraction of municipal solid waste. It is also possible to improve methanogenic communities due to the inoculation of hydrolytic strains. In this regards, Jung (2012) investigated the impact of bioaugmentation with the mesophilic cellulose-degrading strain *Clostridium cellulolyticum H10* on anaerobic digestion of cattle manure and wastewater sludge. This strain breaks down cellulose into hydrogen, acetate, and ethanol, which enhance methanogenesis. One year later, a work from Kovács et al. followed, where they specifically aimed for the inoculation of hydrogenic bacteria to improve methanogenesis. Kovács et al. (2013) investigated the roles of pure hydrogen-producing cultures of *Caldicellulosiruptor saccharolyticus* and *Enterobacter cloacae* in thermophilic and mesophilic natural biogas-producing communities, respectively. Their findings indicated that enhancing biogas production was associated with an increased abundance of hydrogen producers, with the loading rate of total organic solids playing a crucial role in maintaining an altered population balance. Promising results with *Enterobacter cloacae* have again been demonstrated by Ács et al. in 2015.

In another study by Morales-Martínez et al. (2020), the production of hydrogen gas from pretreated agave biomass. Cellulose-degrading microorganisms obtained from bovine ruminal fluid were used to enhance H_2_ production by *Clostridium acetobutylicum*. The results demonstrated the capacity of these microorganisms to hydrolyse the pretreated agave biomass and improve hydrogen gas production, highlighting the potential of bioaugmentation in biohydrogen generation. it was difficult to assign this to a clear phase because H_2_ production relates to acidification but then it was applied in regard to hydrolysis. Collectively, these findings underscore the potential of bioaugmentation strategies in optimising anaerobic digestion processes, promoting higher yields of biogas and hydrogen, and shedding light on the microbial dynamics responsible for enhanced degradation and energy recovery from various organic waste substrates. Such insights can contribute to the development of sustainable and efficient bioenergy production techniques with significant implications for renewable energy applications.

Comparing all 33 articles, species from the following families have been successfully applied in order to improve hydrolysis with the following substrates: *Clostridiaceae* (brewery spent grain), *Thermoanaerobacteraceae* (corn stover), *Thermotogaceae* (sewage sludge), *Flavobacteriaceae* (swine wastewater), *Chlorellaceae* (microalgal biomass), *Fibrobacteraceae* (brewery spent grain) and *Dictyoglomaceae* (cattle manure). As the found strains are mostly related to a better degradation of plant derived material, the importance of lignocellulose needs to be highlighted. It is well known that lignocellulose is the most abundant renewable material on the planet. It is easily accessible and also cost-effective. However, its hydrolysis process is often difficult to complete due to its complex structure.

To move on to the next chapters, the case of Wang et al. (2024) will be described at this point. They evaluated improved thermophilic anaerobic digestion (TAD) of food waste due to four thermophilic strains: *Ureibacillus suwonensis E11*, *Clostridium thermopalmarium HK1*, *Bacillus thermoamylovorans Y25*, and *Caldibacillus thermoamylovorans QK5*. Results showed that cumulative methane production improved by 2.05% (*E11*), 14.54% (*HK1*), 19.79% (*Y25*), and 9.17% (*QK5*) compared to the control. Analysis of microbial community composition revealed increased relative abundance of key hydrolytic bacteria, but also methanogenic archaea. This highlights that the impact on the microbiome cannot just be attributed to the functionality of the strains added. In the particular case of Wang et al., the addition of hydrolytic bacteria was also affecting methanogenic archaea.

### 3.5 Manipulation of acidogenesis

Previous research studies have investigated the role of bioaugmentation in acidogenesis. With the chosen search terms, several articles were found, which were not only addressing anaerobic methane production, but also dark fermentation related articles. Although the present article is primarily not focussed on dark fermentation, these articles were not excluded from the present set of literature. Acidogenesis is a crucial stage responsible for converting complex organic compounds into valuable products such as volatile fatty acids (VFAs) and hydrogen gas. In recent years, microbial bioaugmentation has emerged as a promising approach to enhance acidogenesis efficiency by introducing specific microbial species or mixed cultures into anaerobic systems. This section summarizes and analyses several studies that explore the impact of different microbial bioaugmentation strategies on acidogenesis performance. It needs to be highlighted again that it is difficult to separate between hydrolytic and acidogenic bacteria, because many organisms are able to do both. To cope with this conflict, the authors have shifted articles into the acidogenesis section, if they were discussing the impact on volatile fatty acid (VFA) formation specifically. Articles were also shifted into the acidogenesis section, if an improved hydrogen formation was addressed, but without linking this specifically to syntrophic acetogenesis. Doing this, remaining studies were exclusively related to dark fermentation. In this regard, many of the found articles focus on the production of butyric acid and hydrogen gas (dark fermentation). It needs to be highlighted that production and extraction of butyric acid and/or hydrogen is something, which is usually not wanted in biogas plants, as they are supposed to enrich methane and not hydrogen or organic acids. Therefore, it is crucial to distinguish clearly which of these scenarios (dark fermentation or methane production) is addressed when investigating the impact of bioaugmentation on acidogenesis. To distinguish between hydrolysis and acidogenesis is not always simple. In this regard, Yang et al. (2016) used *L. hydrogenispora ethanolica LX-B* in bioaugmentation experiments, which significantly improved hydrogen production from complex substrates. The bioaugmentation with *LX-B* resulted in hydrogen yields more than twice that of the control group in batch cultivation. Regarding dark fermentation, the improved formation of hydrogen could be subjected to the acidogenesis chapter. However, since improved degradation of complex substrates is addressed, it could also be possible to discuss this article in the hydrolysis section. Another example which fits clearer into the acidification section, has been published by Atasoy & Cetecioglu, (2020). They investigated the enhancement of butyric acid production through bioaugmentation with *Clostridium butyricum* in mixed cultures. Anaerobic sequencing batch reactors were operated under alkaline conditions and fed with dairy industry wastewater as the substrate. Bioaugmentation with *Clostridium butyricum* significantly increased butyric acid production, indicating a positive influence of this specific microbial species on acidogenesis in respect to dark fermentation.

In addition to this, Dams et al. (2016) investigated the potential of bioaugmentation with *Clostridium acetobutylicum ATCC 824* for hydrogen, organic acid, and alcohol production using residual glycerol as the carbon source. Similarily to *Clostridium butyricum*, *Clostridium acetobutylicum ATCC 824* allows for the enrichment of hydrogen. Batch experiments were conducted in pure and mixed cultures, with three different sources of inocula and the experiments were conducted with mixed cultures. The work from Dams et al. is interesting, as it shows the possibility to enrich other metabolites than just hydrogen and butyric acid. Significant yields of hydrogen, but also 1,3-propanediol were achieved when *Clostridium acetobutylicum ATCC 824* was bioaugmented into the sludge from municipal wastewater with 5 g/L of glycerol. One highlight of this work was further the application of glycerol, which is regarded as a recalcitrant substrate. According to Dams et al. and with *Clostridium acetobutylicum ATCC 824* as a microbial additive, glycerol could be a promising substrate for the generation of valuable products like hydrogen and 1,3-propanediol during dark fermentation in mixed culture approaches. Similar as Atasoy & Cetecioglu (2020) or Dams et al. (2016), Goud et al. (2014) evaluated the possibility of bioaugmentation for the improvement of dark fermentation as well. However, they focused their work on indigenous microorganisms, which are naturally present in the environment of dark fermentation processes. They applied three acidogenic bacterial isolates belonging to the phyla *Firmicutes* and *Proteobacteria*, in order to increase hydrogen formation, but also to cope with elevated organic: for this they used the species *Bacillus subtilis*, *Pseudomonas stutzeri*, and *Lysinibacillus fusiformis*. In addition to the work from Dams et al. (2016), Wang et al. (2008) also demonstrated the potential of *Clostridium acetobutylicum ATCC 824* in biohydrogen production through dark fermentation but focused on using microcrystalline cellulose as the carbon source.

Some studies on dark fermentation focus more on the production of organics acids. In this regard, the work by Zheng et al. (2022) can be highlighted. They investigated a biochemical strategy to enhance propionic acid production from kitchen waste acidification through bioaugmentation with *Propionibacterium acidipropionici.* Their results showed that when the inoculum of *Propionibacterium acidipropionici* comprised 30% (w/w) of the seeding sludge, propionic acid production increased by 79.57%.

Most articles that addressed dark fermentation worked with single stages process. In this regard, Liu et al. (2023a) stands out who addressed bioaugmentation in a more holistic approach. They investigated the impact of bioaugmentation technology on anaerobic digestion processes by directly manipulating microbial structure through bioaugmentation in a two-stage co-digestion system. In their study, different doses of *Thermoanaerobacterium thermosaccharolyticum* were introduced into the hydrogen producing pretreatment stage. The system was operating at 55°C. The findings revealed that the addition of *Thermoanaerobacterium thermosaccharolyticum* at 1.12 g had the most significant impact, resulting in cumulative hydrogen and methane yields of 81.54 mL/g VS and 550.98 mL/g VS, respectively. These values were 68.72% and 84.45% higher than those of the control group. Microbial analysis indicated notable changes in microbial community structure, with an increase in the relative abundance of *Thermoanaerobacterium thermosaccharolyticum* during the hydrogen production stage. This increase led to higher levels of volatile fatty acids (VFAs) and hydrogen content, suggesting a potential influence on the acidogenesis step of anaerobic digestion. Like the other acidogenesis related studies, the work from Liu et al. is addressing dark fermentation. However, and unlike the other articles, Liu et al. were the only ones, who implemented this into a system with a subsequent methanation stage.

It needs to be highlighted that bioaugmentation in acidogenesis is not always about improving biomass degradation. It can also be about the elongation of fatty acids. In this regard, a study by Zagrodnik et al. (2020) can be highlighted. They used *Clostridium kluyveri (AS + CK)* in chain elongation processes, where it produced medium-chain fatty acids, such as caproic acid, from a mixed substrate.

### 3.6 Manipulation of acetogenesis

The acetogenesis section explores various strategies to enhance acetate production within AD. Research has focused on utilizing specific microbial strains, such as *Clostridium* and *Thermoanaerobacterium* (Kuribayashi et al., 2017) to improve acetate yields. These strains have shown promise in optimizing the acetogenesis phase by facilitating more efficient conversion of intermediates into acetate. As *Bacillus* species, including *Brevibacillus sp*. *KH_3_* (Li et al., 2009), *Bacillus subtilis* (Xu et al., 2018), and *B. licheniformis* (He et al., 2017), have also been studied for their role in stimulating hydrolytic enzymes and enhancing AD performance, their contributions are more pertinent to the hydrolysis stage and are thus discussed in the corresponding section. In the context of acetogenesis, interactions between acetogens and methanogenic archaea or hydrogen-producing bacteria have been linked to improved biogas production. In this regard, Wang et al. (2018) can be highlighted. They developed a microbial consortium entitled *D83.* For this consortium they highlighted the occurance of *Syntrophospora bryantii*, *Sedimentibacter sp*., and *Thermosyntropha bryantii*, *Methanosarcina sp*. and *Methanobacterium ferruginis*. They further explained that D83 was dominated by hydrogen-producing acetogens, which helped to enhance methane production. Bioaugmentation with *D83* doubled methane yield and rate from glucose fermentation and improved COD removal in molasses wastewater treatment. This study highlighted hydrogen-producing acetogenesis as a key step in methanogenesis, improving both acidogenesis and methanogenesis. There are cases, where such an improvement of methanogenesis is further interwoven with syntrophic relations. Syntrophic bacteria can be involved due to their role in syntrophic acetate oxidation and hydrogen turnover (Zhang et al., 2019). However, the specific role of methanogenic archaea in bioaugmentation is beyond the scope of acetogenesis and is elaborated further in the methanogenesis section. By focusing on acetogenic strains that directly contribute to acetate formation, researchers aim to optimize this critical intermediate step in AD. In a study by Huang et al. (2020), the factors influencing the growth and acetate production efficiency of the *Clostridium sp. NJUST19* strain were investigated under different environmental conditions. The experimental results of digesting waste activated sludge (WAS) with the addition of *Clostridium sp. NJUST19* showed enhanced Total Suspended Solids (TSS) degradation and increased concentrations of VFAs. The TSS degradation rate increased to 35.3%, which was 13.4% higher than the control group. Additionally, the maximum VFAs concentration reached 4200 mg/L, indicating a significant increase of 45.8% compared to the control group. This is another example, which shows how intertwined the different phases of anaerobic digestion are. Although the title from Huang et al. refer to acetogenesis, it remains difficult to differentiate between hydrolysis, acidogenesis and acetogenesis in this specific case. It stands out that in total just one article was fitting into the “acetogenesis” chapter. On one hand, this might indicate a research gap. Amongst the detected articles, there were almost no articles addressing syntrophic butyrate- and propionate-degrading bacteria (SBOBs and SPOBs) and it might be interesting to test the suitability of such organisms for bioaugmentation. On the other hand, missing articles on bioaugmentation with a specific focus on syntrophic, acetogenic bacteria might also be explained by difficulties in culturing such bacteria. It might well be that it is just impractical to use syntrophic, acetogenic bacteria for bioaugmentation. One study that was found, has been published by Shao et al. (2020). They studied bioaugmentation to accelerate recovery in an anaerobic sequencing batch reactor, which was exposed to an organic shock load. The bioaugmented reactor, with a butyric acid-utilizing culture containing *Methanobacteriales* and *Syntrophomonas*, recovered faster (40 days) than the non-bioaugmented reactor (110 days), by relieving feedback inhibition and boosting propionic acid degradation. Another interesting article in the regard comes from Tale et al. (2011), who utilized a propionate-degrading enrichment culture dominated by *Methanospirillum hungatei* and *Methanobacterium beijingense* to bioaugment anaerobic digesters. This approach enhanced recovery after organic overload by reducing acid accumulation and shortening recovery time by approximately 25 days, demonstrating the effectiveness of bioaugmentation in improving process stability. Both studies, the one by Shao et al. and the one by Tale et al., include methanogens, which shows once again that it is not always possible to clearly distinguish published cases regarding the different phases of anaerobic digestion. Yet another case is the study by Akila et al. (2010). They isolated a psychrotrophic xylanolytic acetogenic strain, Clostridium sp. PXYL1, and the *Methanosarcina* strain PMET1 from a cattle manure digester. The addition of PXYL1 increased VFA levels compared to the controls, while the further addition of PMET1 enhanced biogas yields and reduced VFA levels.

An important objective regarding acidogenesis is the improved degradation of long-chain fatty acids (LCFAs). One notable study is by Cavaleiro et al. (2010), which investigated the bioaugmentation of non-acclimated anaerobic granular sludge using *Syntrophomonas zehnderi* to enhance the conversion of LCFAs into methane. In this study, “non-acclimated” refers to the fact that the anaerobic granular sludge had not been previously exposed or adapted to LCFA or similar conditions before the introduction of *Syntrophomonas zehnderi.* The addition of *Syntrophomonas zehnderi* resulted in faster methane production and higher methane yields. In a similar study, Wang et al. (2023) co-cultured *Syntrophomonas wolfei* and *Geobacter sulfurreducens* on the anaerobic anode of a bio-electrochemical system to degrade butyric acid. The co-culture showed a more efficient butyrate degradation than *Syntrophomonas wolfei* and methanogens. The work presented by Wang et al. stands out, as it is not just about adding a certain microbe to the process, but it combines it with the application of electrodes. With their experiment, Wang et al. indicate that the implementation of galvanic elements could be combined with bioaugmentation regarding acidogenesis.

### 3.7 Manipulation of methanogenesis

The stimulation of methanogenesis through bioaugmentation primarily involves enhancing methanogenic archaea, which are crucial for converting hydrogen, carbon dioxide, and acetic acid into methane (Yang et al., 2020). In addition, syntrophic acetate-oxidizing bacteria (SAOBs) are included in this section because they play a unique role in balancing the methanogenesis pathways. Specifically, SAOBs convert acetate into hydrogen and carbon dioxide, thus bridging the gap between acetoclastic methanogenesis (which directly converts acetate to methane) and hydrogenotrophic methanogenesis (which uses hydrogen and carbon dioxide to produce methane). This balancing act helps to optimize the overall methane production in anaerobic digestion processes, making the inclusion of SAOBs in the methanogenesis section particularly relevant. Two pathways can be distinguished for methanogenesis from acetate. The first one is acetoclastic methanogenesis (Zinder & Koch, 1984), in which acetate is enzymatically cleaved into methyl groups (converted directly to CH_4_) and carboxyl groups (oxidised to CO_2_). The second pathway involves a two-step reaction (Zinder & Koch, 1984) performed by so-called syntrophic acetate oxidising bacteria (SAOBs). In this pathway, acetate is first oxidised by SAOBs to H_2_ and CO_2_, and subsequently, these products are further converted to CH_4_ by hydrogenotrophic methanogens.

Due to the syntrophic relationships between methanogens and bacteria, bacteria are relevant for improving methanogenesis, although they usually do not produce methane themselves. In this regard, an interesting work by Zhang et al. (2018) can be highlighted. Zhang et al. was not focussing on methanogens for bioaugmentation. Instead, they were able at low ISR to accelerate methanogenesis by 78% due to the addition of *Geobacter sulfurreducens*. Fluorescence in situ hybridization (FISH) analysis indicated a close association between *Geobacter sulfurreducens* and *methanogens*, which was attributed to syntrophic interactions between *Geobacter sulfurreducens* and methanogens affiliated with *Methanosaetaceae* and *Methanobacteriaceae*. The study from Zhang et al. (2018) is of high interest due to the electrofermentative capabilities of *Geobacter sulfurreducens*. Exoelectrogenic capabilities have the potential to improve syntrophic relations between bacteria and archaea.

Similar to Zhang et al. (2018), a recent work from Zhang et al. (2023) focuses also on bacteria to improve methanogenesis. They used *Lactobacillus lactis* and *Bacillus velezensis* to enhance anaerobic digestion efficiency for food and kitchen waste. Both bioaugmentation and biopretreatment significantly increased crude cellulose removal rates. One might criticise that this finding belongs rather into the hydrolysis section. However, the authors describe that the observed increase in the methane yield (by 22.7–33.6%) was related to an enhanced syntrophic metabolism, which improved hydrogenotrophic methanogenesis. Once again, this shows how deeply intertwined the different phases of anaerobic digestion are. To improve syntrophic relations, it is also possible to apply conductive materials. In this regard, Xiao et al. (2019) stands out, as they explored the combined application of bioaugmentation and conductive materials (CMs). Using *Clostridium pasteurianum* and CMs like biochar and magnetite, they found that hydrogenotrophic and acetoclastic methanogenesis were significantly improved.

Apart from articles focussing on improvement of syntrophic relations, there are many articles the search for way to better cope with ammonium. Ammonia has been identified as a significant inhibitor of methane production in anaerobic digestion. Numerous studies have been conducted to investigate this inhibition and develop strategies to mitigate its adverse effects. In this regard, Yang et al. (2019) presented a work on bioaugmentation, where they tested seven pure strains of microorganisms to recover anaerobic digestion processes, which were suffering from ammonia inhibition. Amongst these strains are obligate acetoclastic methanogens, facultative acetoclastic methanogens, hydrogenotrophic methanogens, and as well syntrophic acetate oxidizing bacteria (SAOBs). The fact that syntrophic bacteria are used to reduce the inhibitory effects of ammonium shows that the two issues are not mutually exclusive. Ammonium thus also seems to have an influence on the syntrophic interaction between bacteria and methanogenic archaea. It is worth noting that obligate acetoclastic methanogens and facultative acetoclastic methanogens are the primary groups capable of converting acetate directly to methane (HAc → CH ₄), which is why they are specifically highlighted in this section. Each of the strains were added to anaerobic digestion processes and the best results were obtained with *Methanobrevibacter smithii* (hydrogenotrophic) co-inoculated with the SAOB *Syntrophaceticus schinkii*. As a result,a 71.1% increase in methane production was observed. Bioaugmentation with *Methanosarcina barkeri* alone proved also to be efficient, enhancing both acetoclastic and hydrogenotrophic methanogenesis with a 59.7% higher methane production. That *Methanosarcina barkeri* showed good results even without the addition of SAOBs can be explained by the fact that *Methanosarcina* can also grow acetoclastically and not just hydrogenotrophically. Interestingly, a negligible improvement was achieved with *Methanothrix*, which is purely acetoclastic. In this regard the work by Chen et al. (2018) is of high interest. They studied the recovery of anaerobic digestion systems under ammonia inhibition. The observed that especially genus *Methanosarcina* recovered fast. The recovery correlated with an increased abundance of the *Firmicutes* genera *Tissierella* and *Lutispora*. Based on these findings, one can argue that both, the hydrogenotrophic and the acetoclastic pathway is important. Yet another study, which confirms the importance of acetoclasic methanogens has been published by Jain et al. (2021), who isolated *T53BJ* from the genus *Methanospirillum spp*. to enhance the biogas yield. Interestingly, Yang et al. also investigated the taxonomic profile upon bioaugmentation. The 16s rRNA gene sequencing results showed that *Methanobacterium spp.* and *Methanothrix spp.* were the dominant archaea in all 14 reactors, regardless of the bioaugmentation. Even after inoculation of *Methanobrevibacter smithii* or *Methanosarcina barkeri*, *Methanothrix* prevailed. In fact, the relative abundances of *Methanobrevibacter smithii* or *Methanosarcina barkeri* remained below <2%. Despite being non-dominant archaea, *Methanobrevibacter spp.* and *Methanosarcina spp.* played pivotal roles in determining the overall microbial consortium and, in turn, improved the overall performance of anaerobic digestion.

In a follow-up study, Yang et al. (2020) used the same strains as mentioned above. This time, they investigated the effectiveness of the respective strains in enhancing methane (CH_4_) production during long-term approaches. The results confirmed the previous findings. Again, *Methanosarcina barkeri* or a combination of *Syntrophaceticus schinkii* and *Methanobrevibacter smithii* resulted in a remarkable increase (35%) in methane potential, although the increase was lower compared to the study before. The lowered increase in the methane potential might indicate that bioaugmentation does not allow a permanent change in the functionality of the underlying microbiome and that bioaugmentation requires a regular and continuous effort of microbial inoculation. Bottles bioaugmented with *Methanosaeta harundinacea (MSH)*, *Syntrophaceticus schinkii*, and *M. smithii* exhibited a significant increment of 49% in methane potential. These findings demonstrate the importance of enhancing both the acetoclastic and hydrogenotrophic methanogenic pathways, while underscoring the need for careful selection of bioaugmentation strains to achieve synergistic effects. Findings as the one from Yang et al. raise the question, whether the inoculation of complex consortia might be more promising than just one strain. Again, Yang et al. performed a taxonomic analysis. Unlike in the study from 2019, *Methanosarcina spp.* prevailed in the archaeal population. Results from Yang et al. are in accordance with a former study from Fotidis et al. (2013). They applied an ammonia-tolerant syntrophic acetate-oxidizing (SAO) co-culture, comprising *Clostridium ultunense spp. nov.* and *Methanoculleus spp*. strain MAB1. This co-culture was tested in a mesophilic up-flow anaerobic sludge blanket (UASB) reactor under high ammonia loads (Fotidis et al., 2013). Bioaugmentation of the SAO co-culture in the UASB reactor alone was not successful, likely due to the slow growth rate of the culture caused by the methanogenic partner. In contrast, when a fast-growing hydrogenotrophic methanogen, *Methanoculleus bourgensis MS2^T^*, was added to the SAO co-culture in fed-batch reactors, a 42% higher growth rate was observed. In accordance with Yang et al., these results show the high potential of using consortia that comprise more than just one strain. Fotidis et al. (2014b) also investigated methanogenic pathways and community composition in full-scale biogas digesters. They investigated these reactors under varying ammonia levels. At high ammonia concentrations, syntrophic acetate oxidation combined with hydrogenotrophic methanogenesis was dominant, with *Methanomicrobiales spp.* (in thermophilic conditions) and *Methanobacteriales spp.* (in mesophilic conditions) being the key methanogens. In contrast, low ammonia levels favoured the acetoclastic methanogenic pathway. Fotidis et al. (2017) presented another study, in which ammonia inhibition was addressed. They have shown that bioaugmentation with ammonia-tolerant *Methanoculleus bourgensis MS2^T^* reversed ammonia inhibition by up to 90% in a continuous stirred tank reactor. Interestingly, Fotidis et al. were not applying a pure culture. Instead, they used an enriched culture, which makes the process more practical. The counteracted ammonia toxicity increased the methane production by 36%. Sequencing revealed a shift in microbial composition, with increased bacterial diversity and reduced archaeal diversity in the bioaugmented reactor. Ammonia-tolerant methanogens dominated, outperforming pure cultures by 25%. Similarly to Fotidis et al., Tian et al. (2019) investigated the importance of *M. bourgensis* regarding ammonia inhibition too. They observed that bioaugmentation of *M. bourgensis* under extreme ammonia conditions (11 g NH_4_^+^-N L^-1^) resulted in an immediate increase of 28% in methane production. The fact that multiple studies highlighted the possibility to overcome ammonia inhibition due to addition of *M. bourgensis*, underscores the importance of this species.

In another study on ammonia inhibition, Yan et al. (2020) assessed the application of bioaugmentation with *Methanoculleus sp. DTU887*. Due to the bioaugmentation, a digester fed with the organic fraction of municipal solid waste showed an increase in the methane yield of 21%. Next to the biogas productivity, Yan et al. were also assessing the concentration of VFAs. They observed a reduction of 10% in VFAs compared to the pre-bioaugmentation period, which indicates a more efficient VFA consumption. Two years later, Yan et al. (2022) continued their research on ammonia-tolerant methanogens. They explored a novel bioaugmentation method, combining gel-immobilized ammonia-tolerant methanogens (biogel) with biochar, to alleviate ammonia inhibition in thermophilic anaerobic systems. Four reactors were subjected to ammonia shocks: one with biogel, one with biochar, one with both, and a control. Results showed that reactors receiving both supplements achieved 100% methane production recovery, while the other configurations showed methane production losses. They described further that reaction with biochar, or the combination of biochar and biogel facilitated the adaptation to higher ammonia levels. It attracts attention that multiple studies tackled ammonia inhibition successful by applying *Methanoculleus* or consortia containing *Methanoculleus* (Fotidis et al., 2013; Fotidis et al., 2017; Yan et al., 2021). In yet another study, Fotidis et al. (2014a) revealed that an increase in methane levels in ammonia-rich environments could be directly linked to the presence of *Methanoculleus*. They introduced a fast-growing hydrogenotrophic methanogen, *Methanoculleus bourgensis MS2^T^*, into a reactor with high ammonia levels, achieving an 31.3% increase in methane production. High-throughput gene sequencing showed a 5-fold rise in *Methanoculleus* spp. abundance after bioaugmentation. Although these results on *Methanoculleus* appear quite promising, there are other hydrogenotrophic methanogens, which could also help to overcome ammonia inhibition. In this regard, Wang et al. (2015) worked with four different hydrogenotrophic methanogens, namely *Methanoculleus bourgensis*, *Methanobacterium congolense, Methanoculleus thermophilus*, and *Methanothermobacter thermautotrophicus.* These hydrogenotrophic methanogens were applied together with two SAOBs, *namely Tepidanaerobacter acetatoxydans* and *Thermacetogenium phaeum*. Under different ammonia concentrations (0.26, 3, 5, and 7 g NH4^+^-N L^-1^), all strains showed the potential to improve the process performance. Yet in another study, Gállego-Bravo et al. (2023) studied enhanced methane production from municipal waste by bioaugmenting a thermophilic anaerobic digestion process with a hydrogenotrophic methanogenic community. Interestingly, they did not work on ammonia inhibition. Instead, the bioaugmentation improved methane yield from the organic fraction of municipal solid waste by 4%. This indicates that bioaugmenting anaerobic digesters with hydrogenotrophic methanogens is useful for more scenarios than just ammonia inhibition. Key microbes involved were the archaeon *Methanoculleus* and bacterial order *MBA08*. To overcome ammonia inhibition the application of oxygen might be interesting too. Although not linked to ammonia inhibition, there is a work the combines the application of oxygen and bioaugmentation. Hua et al. (2022) explored the use of micro-aerobic microbial communities at elevated temperatures. By introducing the methanogens *Methanosarcina acetivorans C2A* and *Methanosaeta thermophila NBRC* 101360, they achieved a significant increase in biogas production of about 44.78%.

So far, most articles about methanogenesis addressed ammonia inhibition or improved syntrophic relations. But there are other stressors that could impair methanogenic communities and, in this regard, it could be interesting to apply methanogens, which can better cope with acidosis. That this is possible was already mentioned earlier with the work by Chen et al., who highlighted the fast recovery of *Methanosarcina*. Another interesting work to cope with acidosis has been presented by Savant et al. (2004). They used the acid-tolerant hydrogenotrophic methanogen *Methanobrevibacter acididurans* to enhance methane production and reduce VFA accumulation in acidic anaerobic digesters. In another study, Li et al. (2021) demonstrated that *Methanosaeta* dominated in anaerobic digestion in oxytetracycline contaminated sludges under acidic conditions with a pH of 4.6 at the first compartment of an anaerobic baffled reactor. In a further study by Town & Dumonceaux (2016), they introduced an acetoclastic consortium into acidified batch digesters, which significantly reduced acetate accumulation and increased methane production. PCR analysis revealed a substantial increase in an acetoclastic methanogens related to *Methanosarcina sp*, which highlights once again the high potential of the genus *Methanosarcina*.

### 3.8 Extraordinary approaches in bioaugmentation research

In table 1, the use of controls has been assessed as described further above. Numbers 1 - 4 were used to indicate whether controls were applied. However, not in all experimental set-ups it is possible or useful to have control experiments, where no cells, autoclaved or non-autoclaved cells are added. Several experiments are not focused on biogas production. Some of them have an unusual set-up, which is designed to evaluate selected strains rather than a complex microbiome. In table 1, such extraordinary cases have been defined as “other”. This concerns for example the case of Arkatkar et al. (2020). The authors assessed coculture conditions for multiple strains based on redox activity, electron transfer rate, columbic efficiency, and internal resistances in a microbial fuel cell, which is very different from typical anaerobic digestion experiments. In such experiments, the aim is not to implement a certain strain into a complex microbiome and therefore, no controls are needed. At least not in the sense as it was analysed in table 1. Nevertheless, such experiments have a certain importance for bioaugmentation. Mapping of microbial interactions can help to define conditions, which might be relevant for bioaugmentation in practice. Another exotic example is from Rinland & Gómez, (2015), who have searched for strains that allow better degradation of onion waste. In this case, multiple strains were isolated from onion waste. However, Rinland and Gómez did not work with complete methanogenic communities. They selected strains that showed good degradation capabilities, and in this case, biogas formation was not a suited criterium to evaluate the experimental success. The authors isolated strains from onion waste at different degradation stages and locations. Growth patterns and carbon source utilisation of the isolates were analysed to identify promising candidates. Among the selected strains, *Bacillus subtilis sp.MB2-62* and *Pseudomonas poae VE-74* demonstrated characteristics making them potential candidates for bioaugmentation or pretreatment in anaerobic digestion processes. As control, they always used a sterilised tube without any active microbes. The work from Rinland & Gómez was taken into account as they were screening for microbes with potential for bioaugmentation, although these strains were not applied in bioaugmentation experiments yet.

Amongst the works, which compared different strains in regard to their biogas formation potential, Jones et al. (2010) was also found as an extraordinary example. They focused on methane generation from nonproductive coal. Coal is a rather unusual substrate for biogas formation, which usually is not degraded. However, the respective production sites contain degradable coal intermediates (geopolymers), for which Jones et al. were highlighting their potential in respect to biogas formation as a potential fuel source. The researchers stimulated methane production using two approaches: biostimulation with nutrient supplementation and bioaugmentation with a consortium of bacteria and methanogens enriched from wetland sediment. The approach differs strongly from typical anaerobic digestion approaches, as they were not starting the experiments with manure from animals or water treatment, which is usually the case in the biogas industry. Apparently, the coal had some intrinsic methanogenic activity, which can be stimulated with nutrients. The biogas formation was even better, if they used a mixed culture from wet-lands. However they did not describe any control, where autoclaved cells were added. So it is difficult to say, which amount of biogas could be attributed to the amount of COD, which was present in the cell mixture added.

Although the focus of the present work was not on microbial fuels cells (MFCs), the used search terms also related to some articles, which used this technology. One work that can be highlighted here is from Arkatkar et al. (2020). They worked with pure cultures, which is usually not regarded as bioaugmentation. However, Arkatkar et al. analysed the coculture behaviour of multiple species. A deeper understanding on how different strains interact and how they might be combined is indeed of interest for better understanding of bioaugmentation. Arkatkar et al. performed coculturing experiments in the anodic chamber of multiple strains, namely *Pseudomonas aeruginosa BR*, *Alcaligenes faecalis SW* and *Escherichia coli EC*. Arkatkar highlighted that coculturing *Pseudomonas aeruginosa BR* with *Alcaligenes faecalis SW* or *Escherichia coli EC* improved the energy generation in both cases. Although the primary goal for Arkatkar et al. was to improve the performance of MFCs and not typical digesters, such experiments can help to define synergies between microorganisms. Although not found with the search terms applied in the systematic search for this study, similar works can be found, when specifically searching for this. For example, a recent work has demonstrated a light driven carbon dioxide reduction to methane by *Methanosarcina barkeri* in an electric syntrophic coculture (Huang et al., 2022b). Bagchi & Behera (2021) investigated the impact of bioaugmentation on microbial fuel cells (MFCs) by introducing *Pseudomonas aeruginosa* into anaerobic sludge (MFC_P_) and comparing its performance to a control MFC seeded with mixed anaerobic sludge (MFC_C_). They also tested an additional MFC with intermittent aeration and bioaugmentation (MFC_P+A_). The results showed that MFC_P+A_ produced significantly more electricity than MFC_P_ alone, with a 4% increase compared to MFC_P_ and a 31% increase compared to MFC_C_. This improvement was due to better organic degradation and more efficient electron transfer in the bioaugmented MFCs. The MFC_P+A_ configuration achieved a coulombic efficiency of 10.4%, which was higher than both MFC_P_ (9.7%) and MFC_C_ (4.41%). These findings are relevant to anaerobic digestion because they demonstrate that bioaugmentation with specific microorganisms, along with intermittent aeration, can enhance the efficiency of electron transfer and electricity generation. This approach can potentially be applied to improve anaerobic digestion processes by optimizing microbial activity and enhancing overall system performance.

Finally and in regard to extraordinary approaches, wastewater treatment should be highlighted. Lin et al. (2018) used *Pseudomonas aeruginosa* for denitrification treatment processes. This process is difficult to relate to any of the four phases of anaerobic digestion.

## 4. Conclusions

In conclusion, the literature underscores the significant potential of augmenting microorganism populations to amplify biomethane production within anaerobic digestion (AD) systems. Bioaugmentation offers the potential to improve yield, speed and robustness through increased biomass conversion, faster digestion rates and / or enhanced process stability. While much of the research has been confined to laboratory settings, the prospect of scaling-up these strategies appears promising. The focus of bioaugmentation primarily on the hydrolysis/acidogenesis phase of anaerobic digestion (AD) is logical, as this stage plays a critical role in enhancing the degradation rate and yield of lignocellulosic compounds. This is particularly important given the abundance of lignocellulosic waste, which serves as the primary feedstock for AD and is often subjected to various stressors. The studies reviewed include a variety of microbial populations, ranging from single species to simple and complex consortia. Notably, multiple articles have shown that the augmentation of just one or a few species can significantly impact the composition of the entire microbiome, underscoring the importance of a metataxonomic approach to study the dynamics of bioaugmentation. Best practice recommended by the authors is to not only map the bioaugmentation culture itself and the microbiome of the anaerobic digester inoculum before bioaugmentation, but also monitor the anaerobic digester after bioaugmentation. This approach helps in understanding how bioaugmentation influences the microbial community structure and functionality over time, which is crucial for optimizing AD processes. Alongside taxonomic screening we recommend putting more attention on conducting the research at a recommended ISR to ensure practical relevance of the results. Moreover, research has consistently demonstrated that the addition of co-cultures or small consortia often produces more significant effects compared to the augmentation of single species. This highlights the potential of mixed-culture bioaugmentation as a promising field for further study, as these consortia can better mimic natural microbial communities, leading to more robust and efficient degradation processes. Among the microorganisms studied, numerous efficient species, such as those from the *Clostridiaceae* family, have shown particular promise in enhancing the hydrolysis/acidogenesis phase. Hydrogenotrophic methanogenic archaea have a great potential in improving digester robustness, it stands out that *Methanoculleus* was used most often. Additionally, despite the few articles, *Methanosarcina* methanogens exhibit remarkable resilience and versatility in biomethane production, making them prime candidates for enhancing the methanogenesis phase and avoiding the accumulation of acetate or hydrogen in digester systems. This improved speed and robustness is opening up the possibility of increasing digester OLR. Despite their potential, research into their utilization in bioaugmentation remains limited. One reason for this might be the difficulties in culturing them in pure culture. Leveraging these archaea could yield substantial benefits, particularly in addressing volatile fatty acid accumulation during the hydrolysis/acidogenesis phase. Thus, further exploration and implementation of bioaugmentation strategies, especially involving mixed cultures and key species like *Methanosarcina*, hold great promise for optimizing AD processes and advancing sustainable biogas production.

## Acknowledgements

The authors gratefully acknowledge financial support from the European Union under the MICRO4BIOGAS project (reference ID 101000470), funded by the European Union’s Horizon 2020 research and innovation programme.

## Author contributions

**MA**: Conceptualization, Methodology, Software, Formal Analysis, Investigation, Resources, Data Curation, Writing – Original Draft, Visualization**; JT**: Conceptualization, Methodology, Writing – Review & Editing**; ML**: Methodology, Software, Writing – Original Draft, Visualization. **CA**: Conceptualization, Methodology, Data Curation, Writing – Review & Editing, Supervision, Project Administration, Funding Acquisition.

## Notes

### Competing Interest Statement

The authors have declared no competing interest.

